# Lytic/Lysogenic Transition as a Life-History Switch

**DOI:** 10.1101/2023.01.03.522622

**Authors:** Joan Roughgarden

## Abstract

The transition between lytic and lysogenic life cycles is the most important feature of the life history of temperate viruses. To explain this transition, an optimal life-history model is offered based a discrete-time formulation of phage/bacteria population dynamics that features infection of bacteria by Poisson sampling of virions from the environment. The time step is the viral latency period. In this model density-dependent viral absorption onto the bacterial surface produces virus/bacteria coexistence and density-dependence in bacterial growth is not needed.

The formula for the transition between lytic and lysogenic phases is termed the “fitness switch”. According to the model, the virus switches from lytic to lysogenic when its population grows faster as prophage than as virions produced by lysis of the infected cells, and conversely for the switch from lysogenic to lytic.

A prophage that benefits the bacterium it infects automatically incurs lower fitness upon exiting the bacterial genome, resulting in its becoming locked into the bacterial genome in what is termed here as a “prophage lock”.

The fitness switch qualitatively predicts the ecogeographic rule that environmental enrichment leads to microbialization with a concomitant increase in lysogeny, fluctuating environmental conditions promote virus-mediated horizontal gene transfer, and prophage-containing bacteria can integrate into the microbiome of a eukaryotic host forming a functionally integrated tripartite holobiont. These predictions accord more with the “Piggyback-the-Winner” hypothesis than with the “Kill-the-Winner” hypothesis in virus ecology.

## Introduction

Viruses have enormous ecological impact (Suttle 2007, Youle *et al*. 2012, O’Malley 2016). Marine viruses infect approximately 10^23^ bacterias every second, removing 20%–40% of the standing stock of marine bacterias each day (Suttle 2007). They are able to exert top-down control of marine microbial abundance comparable to that of zooplankton grazing, thereby affecting marine food webs and biogeochemical cycles (Breitbart *et al*. 2018). Furthermore, viruses represent 94% of the nucleic-acid-containing particles in the oceans (Breitbart *et al*. 2018). Hence, virus also represent an enormous number of evolutionary agents whose phenotypes are shaped by natural selection.

The most important feature of the life history of temperate viruses is their transition between lytic and lysogenic strategies, that is, from a strategy in which a virus infects a bacterium, commandeers its reproductive machinery to produce new viruses and then kills the infected bacterium when releasing the newly minted viruses into the environment *vs*. a strategy of incorporating into the bacteria’s genome as a “prophage” and reproducing along with the bacterium itself. Lytic virus are particularly important for food web dynamics and biogeochemical cycles because “lysis”, that is, the breakdown of the bacterial cell wall, allows the contents of bacterial cells to spill into the media as dissolved organic matter (DOM). As a result, the carbon in infected bacterial cells is intercepted on its journey up the food web—this interception is called the “viral shunt” (Suttle 2007). The viral shunt contrasts with zooplankton grazing on bacteria whereby carbon is passed up the food web when the zooplankton are themselves preyed upon. Thus, which strategy the virus adopts greatly influences its ecological impact. It is lytic viruses that cause bacterial mortality along with the viral shunt, whereas lysogenic viruses do not cause bacterial mortality or a viral shunt. Furthermore, prophages often carry extra genes not pertaining to virus’s own replication but that are expressed by the infected bacterium along with the bacteria’s own genes. These extra genes supplied by the prophage may benefit the bacterium, for example, by endowing it with resistance to antibiotics or by producing pathogenicity to would-be grazers. The virus is a symbiont that can be thought of as a commensal or mutualist when existing as a prophage or as a parasite when existing in the lytic phase. Therefore, determining the evolutionary basis for a switch between lytic and lysogenic life history strategies is of inherent evolutionary interest from life-history and symbiosis perspectives and has great ecological significance as well.

Previous models for the lytic/lysogenic switch (Santillán and Mackey 2004, Ding and Lao 2009, Bednarz *et al*. 2014, Mackey *et al*. 2016, Ch. 7, Sinha *et al*. 2017, Weitz *et al*. 2019, Li *et al*. 2020, Cheong *et al*. 2022, Shivam *et al*. 2022) feature systems of nonlinear differential equations often with time delays, assume mass action collision as the mechanism of viral colonization of the host and postulate density-dependent bacterial population growth.

Instead, this paper introduces a new model that differs from previous virus/bacteria population models in being simpler, boiling down to one equation, and uses different assumptions, namely, viral colonization of the host as a Poisson process rather than by mass-action collisions, density-independent rather than density-dependent bacterial population growth, and it avoids delay-differential equations by taking the viral latency period as the time step in a discrete-time equation analogous to the generation time in population-genetic models.

The new model reveals the conditions for virulent phage and bacteria to coexist as well as the conditions for temperate phage to switch from a lytic to lysogenic life cycle and back again. The mathematical formula for the switch is termed the “fitness switch.” The qualitative implications of the fitness switch for the process of virus-mediated horizontal gene transfer (HGT, transduction), for the formation of three-level holobionts consisting of virus, bacteria and host, as well as for various ecogeographic patterns in viral biogeography are developed.

This paper draws on the approach in ecology pertaining to life-history switches such as those between vegetative growth and flowering in annual plants and between tadpole and adult phases in amphibians *etc*. (*cf*. Schaffer 1983). Models of these developmental switches rely on some equation relating fitness to the time of switching. Then the optimal switch time is computed as that maximizing fitness. Here, the switch is between different life cycles rather than different developmental stages. Still, the idea is the same. The transition between lytic and lysogenic life cycles can be found as the boundary between situations in which one or the other life cycle yields the highest fitness.

The terminology used in the phage/bacteria interaction is summarized in the Supplementary Material. A critique of models using delay-differential equations, mass-action colonization, and density-dependent bacterial population growth is also included in the Supplementary Material. Overall then, the purpose of this paper is first, to explain the lytic/lysogenic switch in terms of optimal life-history theory and second, to introduce a new and simple model for phage/bacteria population dynamics. The life-history theory for the switch is intended to offer qualitative predictions but allows some quantitive predictions too. The new population-dynamic model may be preferable to existing models of phage/bacterial dynamics for reasons detailed below and in the Supplementary Material. Still, the reason for creating this new population-dynamic model is for its use in the optimal life-history theory for the lytic/lysogenic switch.

The model pertains initially to virulent phage that are permanently lytic and then to temperate phage that allow switching between lytic and lysogenic life cycles.

### Virulent Phage: Virus/Bacteria Population Dynamics

Figure 1 (Left) offers a diagram of a virulent virus/bacteria life cycle for a system of equations describing virus and bacteria population dynamics. The model is defined in discrete time with the time step being the viral latency period. A time step begins with the state variables, *N*(*t*) and *V*(*t*) for the number of bacteria and free virus particles (virions) in their environmental source pools at time *t*. Then the virus infect the uninfected bacteria according to a Poisson distribution such that some bacteria remain uninfected, some have only one infection, some have two infections *etc*. (Ellis and Delbrück 1939). Poisson searching for hosts was featured in early parasitoid/host models (Nicholson and Bailey 1935) and more recently for bacterial colonization of hosts in Roughgarden (2018, 2020, 2023).

**Figure 1.**
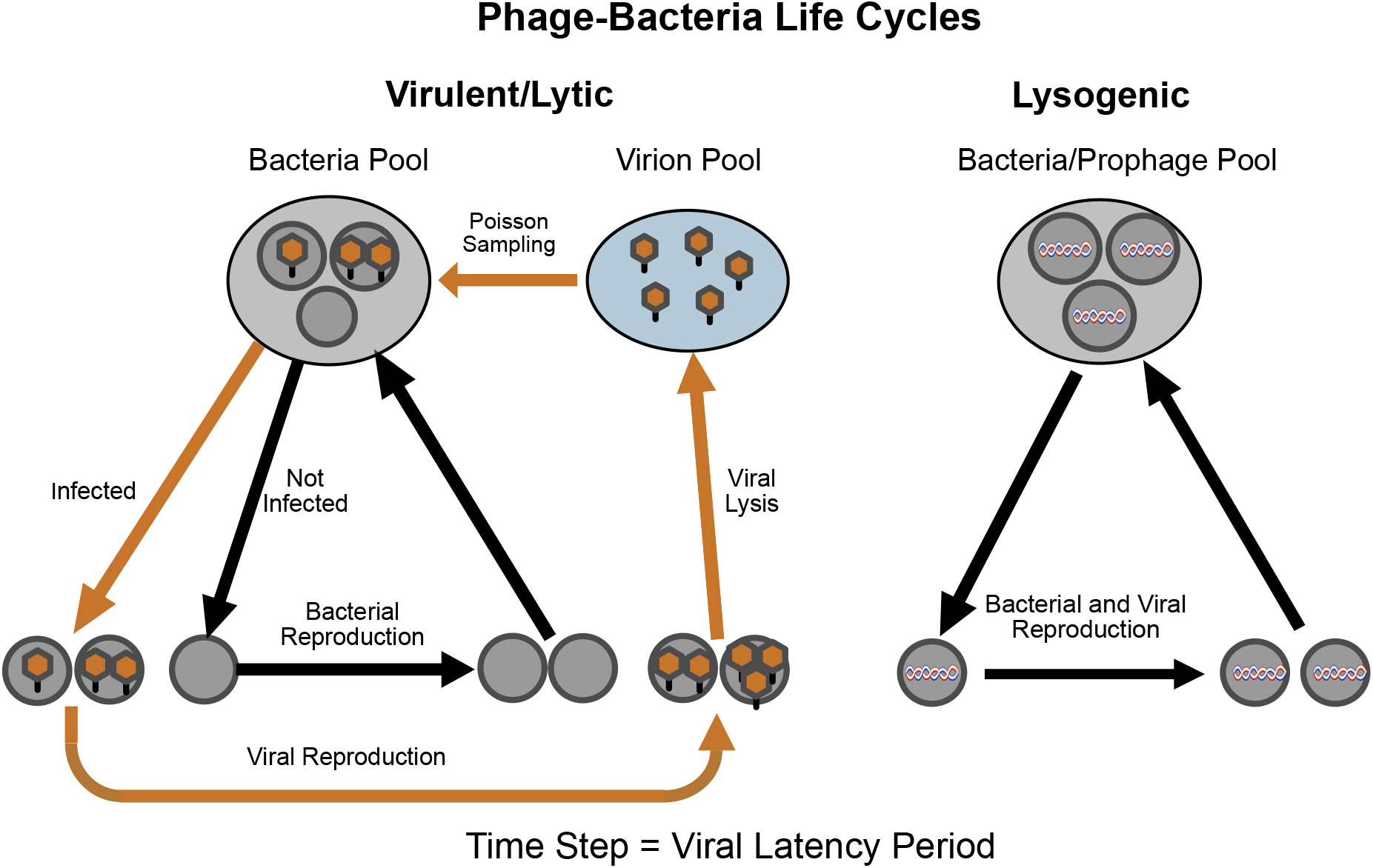
Phage/bacteria life cycles. Left: life cycle for virulent phages and for the lytic phase of temperate phages. Bacteria in the bacterial source pool in the environment are infected by free virus particles (virions) from the viral source pool according Poisson sampling. The infected bacteria are the site of viral replication, leading to the release of new virions and the death of the infected cells. The uninfected bacteria replicate to produce more bacteria available for infection to begin the next time step. Right: life cycle for the lysogenic phase of temperate phages. The virus is represented as a helix that is incorporated as a prophage into the genome of its bacterial host. The virus life cycle is coincident with the bacterial life cycle. The time step is the viral latency period.

The Poisson distribution describing the infection process has density parameter, *μ*(*t*) *≡ a*(*v*(*t*)) *v*(*t*) where *v*(*t*) *≡ V*(*t*)/*N*(*t*) is the ratio of virus to bacteria in their source pools. The *a*(*v*(*t*)) is the adsorption function—it refers to the probability that a viral particle comes in contact with a bacterium and binds to surface molecules for subsequent incorporation into the bacterial cell. Adsorption is a function of *v*(*t*) because the number binding sites on the surface of the bacterium is finite and the sites may become saturated. Thus, *a*(*v*(*t*)) includes information about the binding affinity of virus particles to receptors in the bacterial surface and also about the number of binding sites. Furthermore, *a*(*v*(*t*)) also includes information about the accessibility of the bacterium to virus particles and has a role similar to the colonization parameter, *d* in a model for bacteria colonization of a host to form a holobiont (Roughgarden 2023). For a given ratio of virus to bacteria, *v*(*t*), if the environment is such that virus are prevented from reaching the bacteria, then *a*(*v*(*t*)) is low compared to *a*(*v*(*t*)) in environments that allow ready access of the virus to the bacteria. Environments may differ in how viscous the media is, or for airborne virus particles, whether air filters are installed.

### Recursion Equations in Discrete Time

According to a Poisson distribution, the fraction of uninfected bacteria that remain uninfected, *P*_0_ (*μ*(*t*)), is *e*^*−μ*(*t*)^ and the fraction of bacteria colonized by *m* virus, *P*_m_ (*μ*(*t*)), is 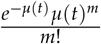 for *m ≥* 0 where *μ*(*t*) = *a*(*v*(*t*)) *v*(*t*). The *m* indexes the number of virus per bacterium. The fraction of bacteria that are infected by one or more virus is 1 *− P*_0_(*μ*(*t*)).

The uninfected bacteria population increases by a geometric factor of increase per time step, *R*. Necessarily, *R >* 1, for the bacteria to be able to increase in the absence of the virus. Accordingly, the uninfected bacteria resupply the bacterial source pool as

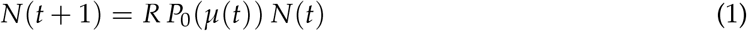

where *P*_0_(*μ*(*t*)) is the Poisson probability that a bacterium is uninfected.

Next, the virus in the infected bacteria produce new virus particles as

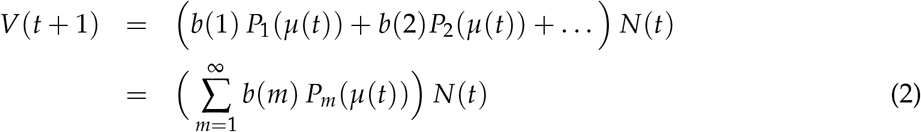

where *b*(*m*) is the burst size from a bacterium infected with *m* virus particles and *P*_*m*_(*μ*(*t*)) is the Poisson probability that a bacterium is infected by *m* virus particles. More succinctly,

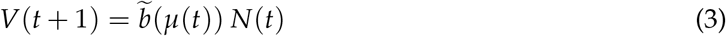

where

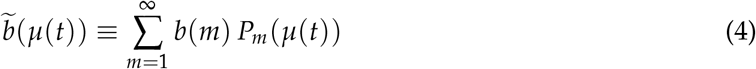

and 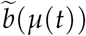 (*μ*(*t*)) is the burst per bacterium from all the infected bacteria at time *t*.

Eqs. 1 and 2 together constitute a population-dynamic model for virulent virus and bacteria populations based on the life cycle sketched in Figure 1 (Left). The parameters in the model are *R*, the geometric factor of increase for uninfected bacteria, *a*(*v*), the adsorption function, and *b*(*m*), the burst-size function.

The model may be simplified by dividing Eq. 3 by Eq. 1

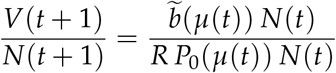

Then identify *v*(*t* + 1) with 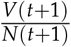 and *v*(*t*) with 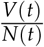 yielding

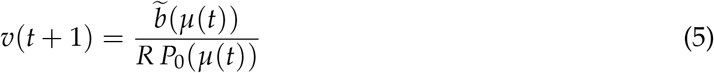

Equation 5 is a dynamical equation in one variable, *v*(*t*), where *μ*(*t*) = *a*(*v*(*t*)) *v*(*t*).

Density dependence in the burst size, *b*(*m*), and/or the absorption function, *a*(*v*), potentially leads to a stable equilibrium viral to bacteria ratio, 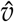. The 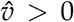 is a necessary condition for stable coexistence of a virulent virus and its host. Alternatively, for some parameter values, 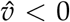, indicating that the virus cannot coexist with the host. Still another possibility is that the virus can coexist with the host, 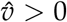, but the virus’ impact on the host is sufficient to drive the bacteria (and itself) to extinction.

This model does not require an assumption of density-dependent bacterial population growth. The stability, or lack thereof, of virus/bacteria coexistence in this model is determined solely by properties of the virus/bacteria interaction itself as contained in the *b*(*m*) and/or *a*(*v*) functions. That is to say, the model may be thought of as pertaining to the log phase of bacterial population growth. Once a stable virus/bacteria equilibrium ratio is reached during the bacterial log phase, both parties possess a common geometric growth factor. Then, external to the virus/bacteria interaction, density dependent might *also* occur in the bacterial population dynamics, *e.g*., the bacteria might grow according to a logistic equation. If so, after attaining coexistence at a stable virus/bacteria ratio during their log phase, the population pair can proceed to grow together with their common geometric growth factor eventually reaching a joint stationary phase determined in part by the microbial carrying capacity.

This qualitative picture of virus/bacteria population dynamics is fundamentally different from the classical picture proposed in Campbell (1961) and later. The Campbell model envisions that virus/bacteria population dynamics involves two counteracting forces—the stabilizing force of density-dependent bacterial population growth and the destabilizing force of time lags arising from the virus latency period. The outcome is a balance of these forces. Accordingly, the formulas for the virus/bacteria equilibrium and its stability in the Campbell model depend on value of the microbial carrying capacity as well as on parameters pertaining to the virus/bacteria interaction. This commingling of parameters for the virus/bacteria interaction with those for bacterial population growth leads to peculiar predictions.

For example, according to the formula for the equilibrium in the Campbell model (*cf*. Eq. S6), small mammals should have fewer phage infecting bacteria than large mammals because the low bacterial carrying capacity in their small guts precludes the coexistence of many types of phage. Conversely, large mammals should support more phage infecting their bacteria because the large carrying capacity for bacteria in their large guts allows more types of phage to coexist with bacteria. However, this phage/bacteria coexistence in large mammals may be temporary because the weakened density dependence from a large bacterial carrying capacity should lead to large virus/bacteria oscillations. No evidence supports this qualitative picture.

The question of which qualitative picture applies to the phage/bacteria interaction, namely that offered by the present model *vs*. that offered by the classical model of Campbell (1961), will ultimately need to be settled empirically. In the meantime, a comparison of the models might be assisted by the Supplementary Material that offers a detailed mathematical summary of the Campbell (1961) model as well as discussion of its quantitative and technical limitations.

Turning now to the present model, density dependence in burst size might, in principle, underwrite stable coexistence of virus and bacteria. However, the evidence shows that density dependence in burst size is weak (*cf*. Gadagkar and Gopinathan 1980, Figs. 4–5, Patel and Rao 1984, Fig. 1–2). Using these published data relating burst size to *m*, analysis in the Supplementary Material shows that density dependence in burst size does not allow for stable virus/bacterial coexistence. In contrast, it will be shown here that density dependence in viral absorption onto the bacterial surface does allow stable virus/bacteria coexistence.

**Figure 2.**
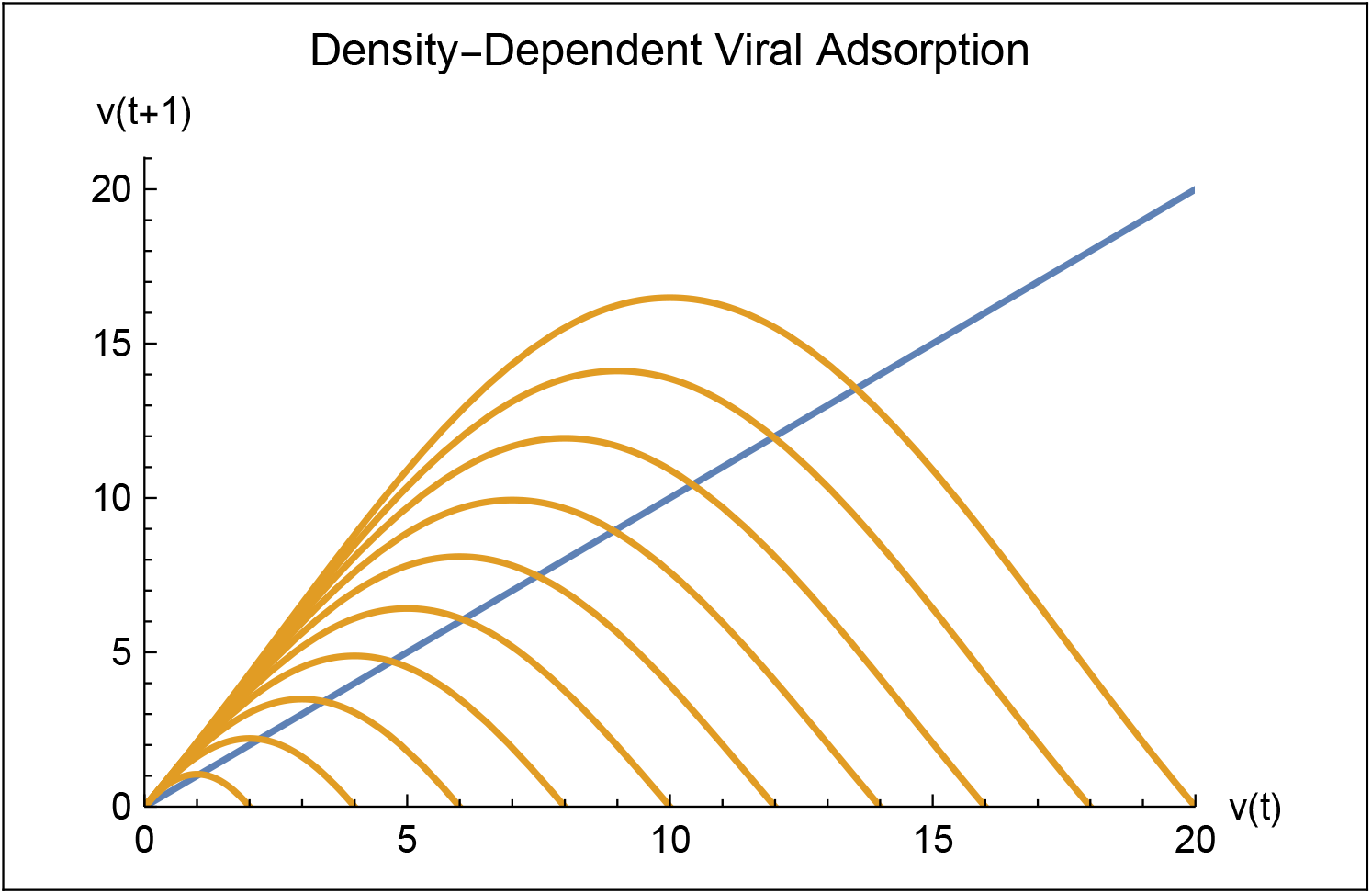
v(t + 1) vs. v(t), in brown, for density-dependent absorption from Eq. 8. The diagonal line, in blue, indicates v(t + 1) = v(t). The intersection of the brown curve with the blue line indicates an equilibrium point where virus and bacteria coexist provided the equilibrium is stable. Here, a_o_ = 0.1, b_o_ = 40, R = 2, and κ varies left to right from 2 to 20 in steps of 2.

The molecular biology of viral absorption onto the surface of bacterial and animal cells has been the focus of much recent study (Dimitrov 2004, Grove and Marsh 2011, Yamauchi and Helenius 2013, Maginnis 2018, Koehler *et al*. 2020, Huss *et al*. 2021). The kinetics of virus binding to cell surface has also been investigated (Moldovan *et al*. 2007, Hubbs *et al*. 2019, Echeverría-Vega *et al*. 2020, Koonjan *et al*. 2022). These studies do not present data directly on *a*(*v*) but are nonetheless consistent with the assumption that the binding sites for virus on the surface of a bacterium become saturated as the density of viral particles increase. Accordingly, the absorption function will be taken as 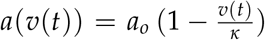 where *κ* is the saturation density—the viral density at which all the binding sites per bacterium are occupied and *a*_*o*_ is the maximal probability of binding when all the binding sites are available, given the environmental conditions. Also, it is required that 0 *≤ v*(*t*) *≤ κ*. This formula postulates that the probability of a viral particle binding to a bacterium tends to zero as the binding sizes fill up, and equals zero when the viral density, *v*, equals *κ*.

Because density-dependent absorption does allow stable virus/bacteria coexistence, subsequent analysis assumes solely density-dependent absorption without any density dependence in burst size.

**Density-Dependent Absorption:** 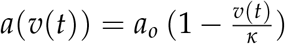, *b*(*m*) = *b*_*o*_ *m*

Here,

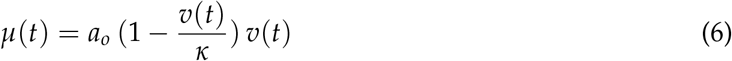

and, upon doing the summation in Eq. 4^3^,

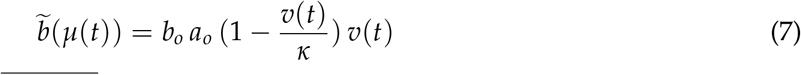

so Eq. 5 becomes

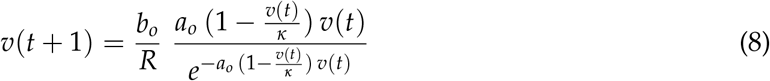

Illustrative parameter values roughly follow those used in the Supplementary Material for the Campbell (1961) model. The time step, which is the latency period, is 15 minutes (Endy *et al*. 1997, Purohit *et al*. 2005), implying four steps per hour. Using the Campbell value for *r* = 0.0347 for an *E. coli* doubling time of 20 minutes leads to a geometric factor of bacterial population growth per time step, *R*, of 2.68 *i.e*. (1 + *e*^0.0347*×*15^). For simplicity, *R* is taken as 2. The per-virus burst size, *b*_*o*_, is taken as 40 following Gadagkar and Gopinathan (1980, Figure 4). The collision rate, *c*, in the Campbell model was taken as 10^*−*4^ following Philipson (1983). This collision rate per binding site times the number of binding sites, estimated as 10^3^ in Philipson (1983), leads to an initial viral absorption coefficient, *a*_*o*_ as 0.1 For the density-dependent absorption, *κ* is arbitrarily taken as 2 to 20 with a increment of 2. These parameters appear in Table 1.

**Table 1:** Example Parameters in Model.

|  |  |
| --- | --- |
| Time Step | 15 min |
| $R$ | 2 |
| $b_o$ | 40 |
| $a_o$ | 0.1 |
| $\kappa$ | 2 to 20, Increment 2 |

Figure 2 shows the curve for *v*(*t* + 1) from Eq. 8 and the line for *v*(*t* + 1) = *v*(*t*) for several values of the saturation density *κ* based on the parameter values in Table 1. The curves intersects the line indicating equilibrium points, 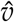, at which virus and bacteria can coexist, providing the equilibrium points are stable, as analyzed further in the next sections. The equilibrium points, 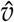, increase with increasing saturation densities, *κ*. Moreover, the slope of the *v*(*t* + 1) curve at the intersection point is always negative and become steeper as *κ* increases.

### Condition for Viral Increase when Rare

The virus population can begin to infect the bacterial population if it can increase when rare. That is, if *v*(*t*) *≈* 0, then *v*(*t* + 1) *> v*(*t*) if

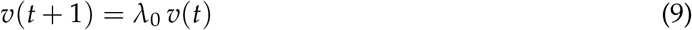

where

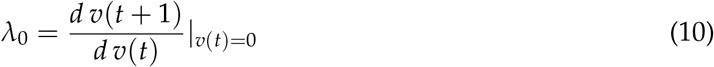

The *λ*_0_ works out to be

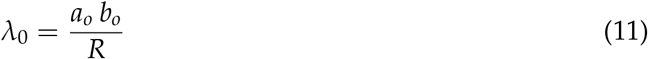

The *λ*_0_ must be greater than 1 for the virus to increase when rare, according to Eq. 9. Now, *R >* 1 is the condition for the bacteria to increase in the absence of the virus, which is greater than one by assumption. Therefore, if *a*_*o*_ *b*_*o*_ *> R >* 1 the virus can spread into a population of increasing bacteria. In words, this condition means that *the initial absorption probability times the per-virus burst size, which is a viron’s expected production per time step, must exceed the bacterium’s production per time step for the virus to spread when rare. Otherwise, the bacteria population outgrows the virus population—the virus cannot keep up*.

Given *b*_*o*_, and *R*, then Eq. 11 implies that the absorption probability must exceed a minimum for the virus to spread. Putting *λ*_0_ = 1 and solving for *a*_*o*_, yields the minimum probability of absorption that permits the virus to spread,

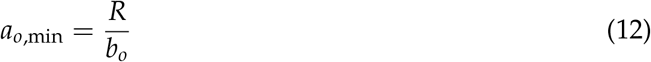

In the instance of the parameters in Table 1 where *b*_*o*_ is 40 and *R* is 2, *a*_*o*,min_ works out to be 0.05. Hence *a*_*o*_, which has been taken as 0.1, is above the minimum. Given the burst size and the bacterial growth factor, if the absorption probability is less than *a*_*o*,min_ then the virus cannot spread into a growing population of bacteria. Environmental interventions to halt the spread of the virus must reduce *a*_*o*_ below *a*_*o*,min_ to succeed.

### Equilibrium Viral Density, 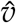

Given that both the virus and bacteria can increase when rare, they may come to the equilibrium at an intersection between the *v*(*t* + 1) curve and the *v*(*t* + 1) = *v*(*t*) line, as was illustrated in Figure 2. The point of intersection depends on the saturation density, *κ*. The **FindRoot[]** function in *Mathematica* is used to calculate the value of 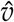 at an intersection point. Figure 3 (left) shows the curve of 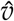 obtain in this way as a function of *κ*.

**Figure 3.**
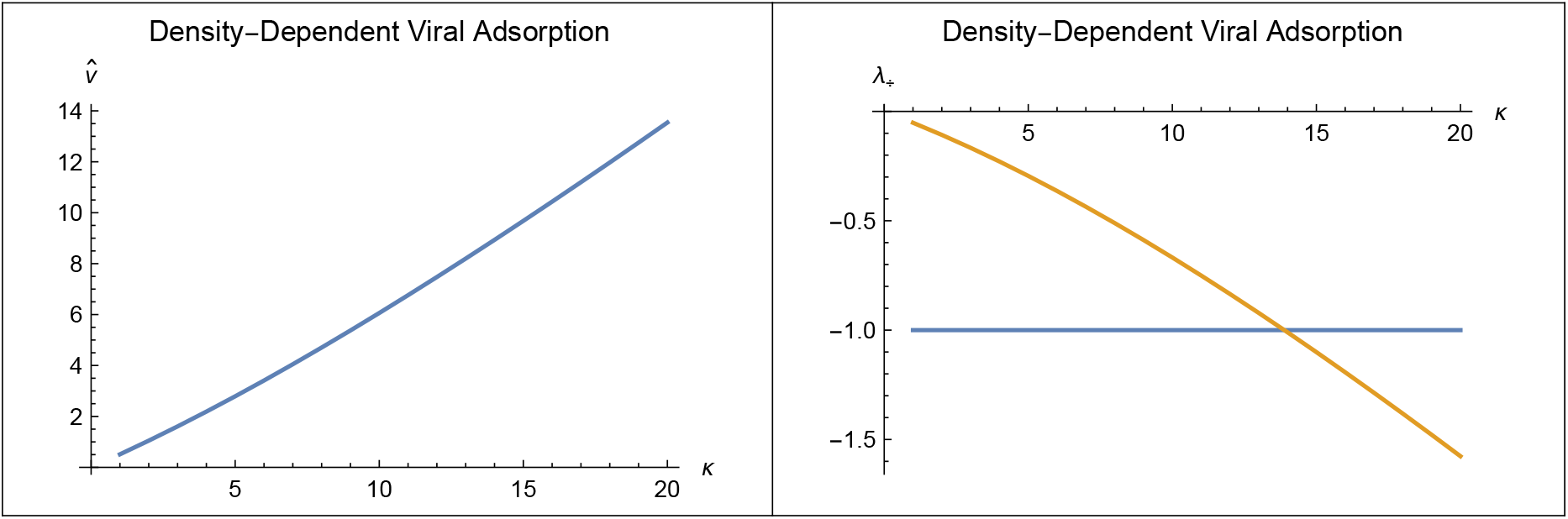
Left: Equilibrium viral ratio, 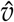 as a function of κ. Right: Curve is eigenvalue evaluated at 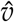 as a function of κ and line at −1 marks the boundary between stable and unstable equilibria. The equilibria for κ below 14 are stable and characterized by a damped oscillatory approach whereas the equilibria for κ above 14 are unstable. Here, a_o_ = 0.1, b_o_ = 40, R = 2, and κ varies from 1 to 20.

**Figure 4.**
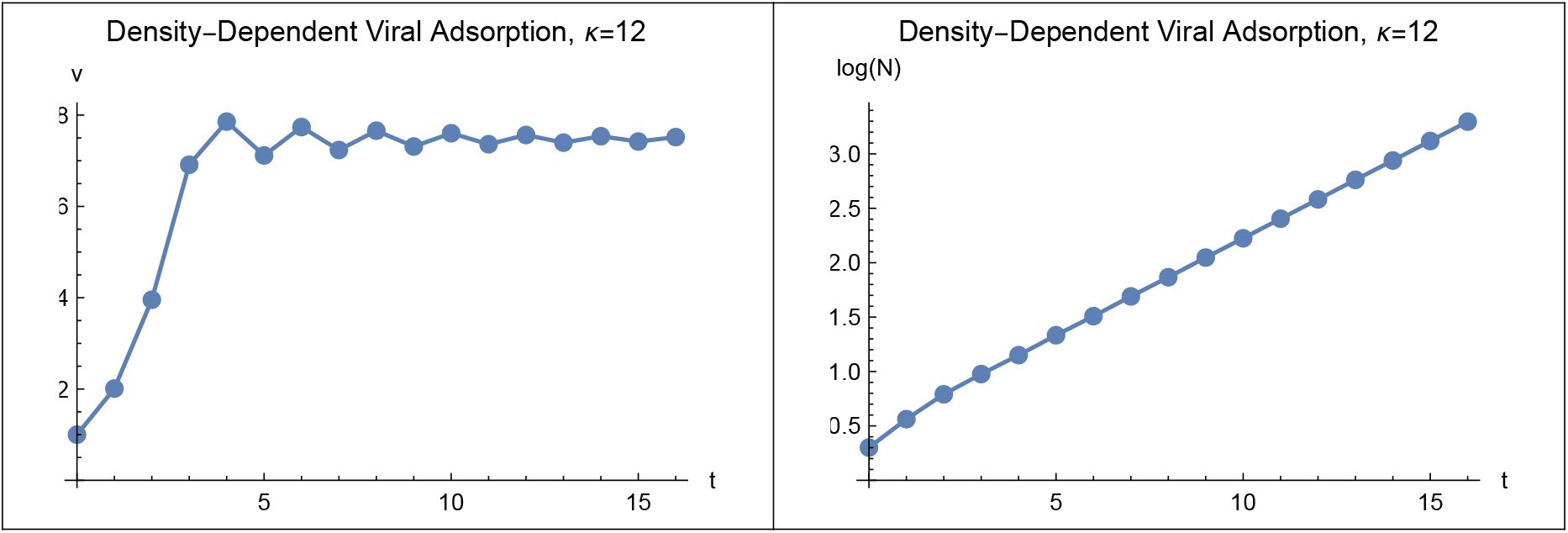
Left: Trajectory of v(t) showing damped oscillation during approach to equilibrium. Right: Trajectory of log_10_(N(t)) showing approach to exponential bacterial population growth. Here, κ = 12, v(0) = 1, N(0) = 2, t_max_ = 16, κ = 12, a_o_ = 0.1, b_o_ = 40, R = 2 and t_max_ = 16.

**Figure 5.**
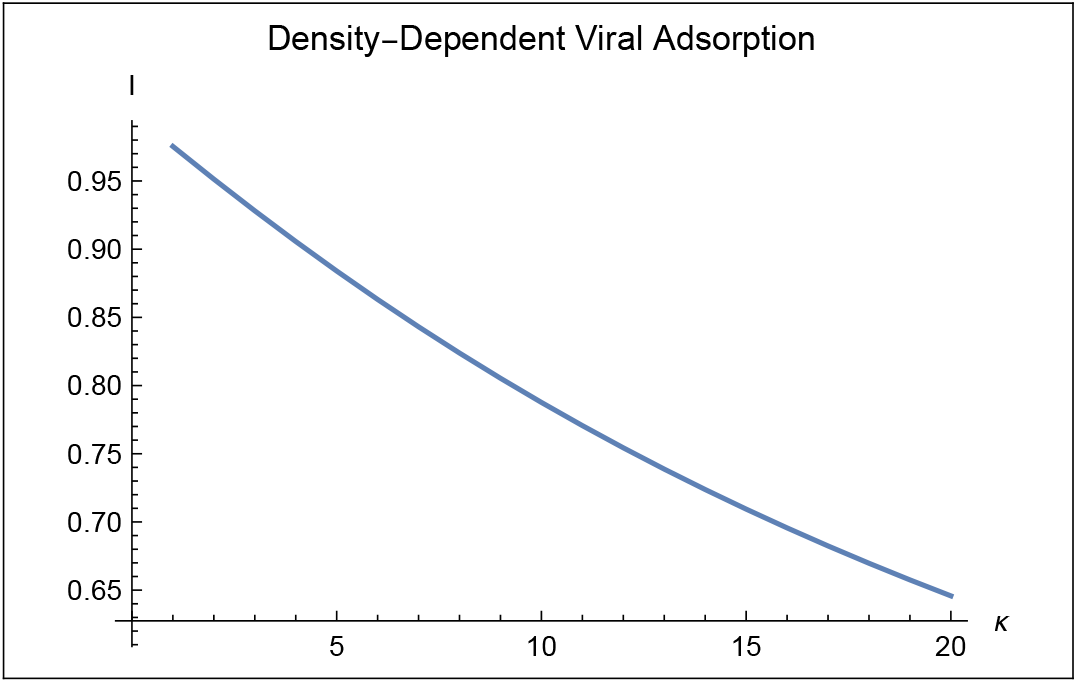
Infection discount. Here, a_o_ = 0.1, b_o_ = 40, R = 2 and κ varies from 1 to 20.

### Stability of 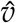

The stability of an equilibrium viral ratio, 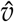, depends on *λ*_+_ defined as

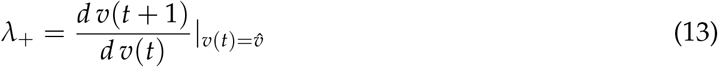

The *λ*_+_ is the eigenvalue evaluated at a positive equilibrium point. If *λ*_+_ is between +1 and *−*1 then the equilibrium is stable (*e.g*. Roughgarden 1996). Specifically, if *λ*_+_ is between +1 and 0, the approach to equilibrium does not involve oscillations whereas if *λ*_+_ is between *−*1 and 0, the approach to equilibrium involves damped oscillations.

Figure 3 (right) shows the curve of *λ*_+_ as a function of *κ*. The *λ*_+_ is always negative, so the approach to equilibrium is oscillatory. The horizontal line at *−*1 marks the boundary between stable and unstable equilibria. The horizontal line crosses the curve for *λ*_+_ near *κ* = 14. Hence, the equilibria, 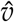, for *κ <* 14 are stable with an oscillatory approach. The equilibria for *κ >* 14 are unstable.

An additional source of density dependence may smooth the approach to equilibrium. Indeed, if density dependence in burst size is combined with the density dependence in absorption, then the eigenvalue may lie between 0 and +1, indicating a non-oscillatory approach to the equilibrium (not shown).

### Trajectories

Figure 4 illustrates trajectories of both virus and bacteria for *κ <* 14. Trajectories illustrate the damped oscillations of *v*(*t*) during the approach to a stable equilibrium. The trajectory of *N*(*t*) also shows damped oscillations during the approach to the asymptotic growth factor, although the scale on the graph makes the oscillations difficult to see. The Supplementary Material also includes an illustration of trajectories around an unstable equilibrium where *κ >* 14.

### Infection Discount

Figure 4 shows the bacterial population increasing with time. However, unlike this figure, the bacterial population does not necessarily increase even though the virus to bacteria ratio has reached a stable equilibrium, 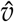.

The net growth factor, *R*^*′*^, for the bacteria per time step from Eq. 9 is *R*^*′*^ *≡ R e*^*−μ*(*t*)^. The *R*^*′*^ must be *>* 1 for the bacterial population to increase. If the virus to bacteria ratio is at equilibrium, 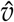, then 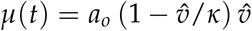. In this situation,

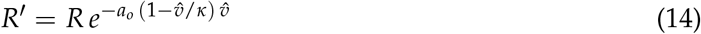

Define the “infection discount”, *I*(*R*), as

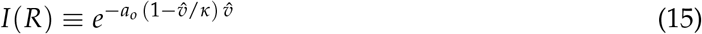

The infection discount indicates the degree to which the bacterial population growth factor is discounted as a result of the mortality caused by viral infections. The infection discount depends implicitly on *R* because 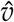 depends on *R* along with the other parameters. Hence, the bacteria can increase if

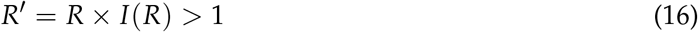

If 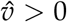 and *R*^*′*^*<* 1 then the virus can increase when rare but then subsequently drives the host (and itself) to extinction.

Figure 5 illustrates the infection discount for the parameters of Table 1. The discount becomes more severe as *κ* increases because 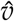 increases with *κ*. However, in the table, *R* is 2 and the discount never drops below 1/2, so *R*^*′*^ remains greater than one even as *κ* increases to 20. This indicates that the bacterial population can continue to increase for all levels of infection with the parameters of Table 1. Other parameters values (not explored) might not be so kind to the bacterial population.

### Temperate Phage: Criterion for Lytic to Lysogenic Transition

The model for the population dynamics of virulent phage in Figure 1 (Left) also applies to the lytic phase of a temperate phage. For the lysogenic phase, a virus incorporates into a bacterium’s genome and reproduces along with the bacterium itself as shown in Figure 1 (Right). The main life-history question pertaining to temperate phage is when to transition from lytic to lysogenic and *vice versa*. The premise of this paper is that a phage should switch in the direction that yields it the highest fitness.

The quantitative answer to the lytic/lysogenic life history question follows directly from the condition for virus to increase when rare, Eq. 11. The population dynamics of lysogenic phase is identically the same as that for the bacteria they reside in, namely, increasing by a factor of *R* each time step. On the other hand, the lytic phase can spread into the population of bacteria if *a*_*o*_ *b*_*o*_ *> R*, meaning that the initial absorption probability times the per-virus burst size exceeds the bacterial growth factor. So the lytic phase produces more virus per time step if if *a*_*o*_ *b*_*o*_ *> R* whereas the lysogenic phase produces more virus per time step if *a*_*o*_ *b*_*o*_ *< R*. Thus, a “fitness switch” can be formulated as follows:

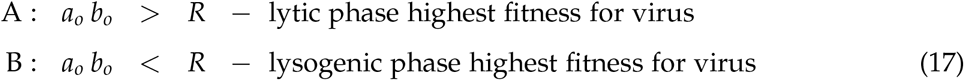

Bruce *et al*. 2021 employ a similar criterion in a four-equation model when comparing the reproductive value of prophage to the reproductive value of lytic virus. Virus communication systems might supply virus with the information needed to activate the switch in one direction or the other (Erez *et al*. 2017, Trinh *et al*. 2017, Harms and Diard 2019, Rajaure and Adhya 2019, Silpe and Bassler 2019, Tan *et al*. 2020, Brady *et al*. 2021, Bruce *et al*. 2021).

Now consider qualitative predictions of Eq. 17.

### Qualitative Predictions of Fitness Switch

#### Laboratory Conditions

The fitness switch agrees qualitatively with well-known laboratory conditions that cause a lytic/-lysogenic switch. Percent lysogeny in a bacterial host is an increasing function of the number of coinfecting virus (Fry 1959, Kourilsky 1973, Kourilsky and Knapp 1974). The interpretation of this finding according to the fitness switch pertains to competition among the coinfecting virus for the cell’s resources. Experimental conditions maintain a constant base absorption, *a*_*o*_, and bacterial growth factor, *R*, among treatments but vary in the number of introduced phages. Only the Poisson density, *μ* = *a*_*o*_ *v*, varies among treatments—this is what leads to varying coinfection among treatments. Furthermore, according to Kourilsky (1973), treatments that depress host macromolecular syntheses favor lysogeny. Hence, one may hypothesize that competition among multiple virus per bacterium affects the number of virons each produces given a limited amount of resources for macromolecular synthesis, thereby lowering the per-virus burst size and making lysogeny adaptive compared with lysis. Even strong density dependence in burst size is not sufficient to stabilize virus/bacteria population dynamics (*cf*. Eq. S17) but density dependent burst size remains relevant to whether an individual virus’ fitness is higher *via* the lytic or the lysogenic life cycle. A similar interpretation has been made by Attrill *et al*. (2023).

The molecular mechanism underlying an adaptive increase in lysogeny with increasing coinfection might involve an increase in copies of phage repressors of the lytic cycle and of integrase genes needed for entry into the host genome (Silveira *et al*. 2021). This mechanism for attaining increased lysogeny with increased coinfection could be considered as bi-product cooperation among the coinfecting viruses. Active cooperation would be indicated if phage repressor and integrase concentrations exceed what is predicted if these concentrations were to scale linearly with the number of coinfections (*cf*. Kourilsky 1973, Trinh *et al*. 2017).

Conversely, damage to the bacterial DNA, say with ultraviolet iradiation, triggers a so-called SOS response within the bacterium that in turn provokes a prophage to revert to a lytic state, a process called “induction” (Bæk *et al*. 2003). Damage to the bacterial DNA would lead to a lower bacterial *R*, making lysis adaptive compared with lysogeny.

Beyond the laboratory, the fitness switch is relevant to various environmental scenarios that flip the switch from lytic to lysogenic or *vice versa*.

#### Intervention Affecting *a*_*o*_ with Hygienic Practice

If the access of virus particles to bacteria is sufficiently lowered by introducing air filters or other hygienic practices, then the best strategy switches from lytic to lysogenic. For example, the minimum value of *a*_*o*_ for the virus to spread in a bacteria population from Eq. 12 using the parameters in Table 1 works out to be 0.05. So if the hygienic practices reduce *a*_*o*_ below 0.05, the the virus should switch from lytic to lysogenic because doing so yields a higher viral population growth. This conclusion is consistent with the conclusion from Stewart and Levin (1984) that “lysogeny is an adaptation for phage to maintain their populations in ‘hard times’.” In this scenario, the virus are using the bacteria as a refuge.

#### Environmental Deterioration Affecting *b*_*o*_ and *R* Simultaneously

If a chemical such as a general toxin, pollutant or antibiotic is introduced into the environment, then whether the virus should switch from lytic to lysogenic depends on the relative impact of the chemical. For example, if the chemical is an antibiotic, then the bacterial growth factor, *R*, is lowered, say by 50%. Now, the per-virus burst size, *b*_*o*_ is likely affected too because, say, the ATP level and other resources within the bacteria drop, implying that fewer viral particles can be synthesized. If the per-virus burst size is also reduced by 50% as a result, the reductions to both virus and bacteria cancel out in the switch function, Eq. 17, leaving the lytic *vs*. lysogenic state unchanged. But if the chemical sufficiently affects *b*_*o*_ more than *R*, then the virus could switch from lytic to lysogenic and conversely, if the chemical affects *R* more than *b*_*o*_. Thus, the idea of lysygeny offering a refuge for lytic virus to escape “hard times” depends on the relative impact of the deterioration on the virus and on the bacteria.

#### Viral Biogeography of Lysogeny and Pathogenicity

Several correlations among habitats in the propensity for lysogeny have been discovered. Here is how these patterns might be accounted for based on the fitness switch, Eq. 17.

#### Microbialization on Coral Reefs—Habitat with High *R*

Overfishing and eutrophication on coral reefs leads to an increase in fleshy algae. The algae are largely ungrazed and produce an increase in dissolved organic carbon (DOC). The increase in DOC in turn leads to a replacement of the type of bacteria characteristic of low-nutrient environments (oligotrophs) with those typical of nutrient rich environments (copiotrophs). Forest Rohwer and colleagues (Haas *et al*. 2016) refer to this shift in coral reef ecosystem structure as “microbialization.” Furthermore, based on 24 coral reefs, they report (Knowles *et al*. 2016) that viromes in microbialized conditions show an increase in the marker genes of temperate viruses, indicating that increased lysogeny accompanies microbialization.

The increase in DOC from microbialization leads to an increase in microbial growth, *R*. The effect of increasing *R* in the switch function, Eq. 17, provided that *b*_*o*_ is relatively unchanged, is to switch the virus from lytic to lysogenic, *i.e*. from state A to state B. The fitness switch therefore can predict that the microbialization resulting from increased DOC leads to an increase in lysgenic bacteria, as has been observed.

Rower and colleagues (Haas *et al*. 2016, Silveira *et al*. 2020) further observe that the enrichment of copiotrophs during microbialization is correlated with an increase in virulence genes (Vega Thurber *et al*. 2008). This in turn is correlated with an increase in the prevalence of coral disease (Harvall *et al*. 1999). They conjecture that the pathogenicity of copiotrophs compared with that of oligotriophs is caused by prophages accumulating in the copiotrophs that carry virulence genes for protection of their bacterial hosts. Expression by the bacteria of the virulence genes carried by the prophages produces virulence factors (Bondy-Denomy and Davidson 2014, Davies *et al*. 2016, Obeng *et al*. 2016).

The virulence factors from these genes are of no *direct* value to the virus and its replication within the host, but by assumption, offer protection to their bacterial host. If so, the bacterial *R* increases because of this increased protection. This increase in *R* due to the virulence genes being expressed adds to the increase in *R* caused by the increased DOH that causes microbialization to begin with.

The population growth of the virus as a prophage is identically the same as that for the bacterium, *R*. So, any assistance the prophage supplies to its bacterial host that increases the bacteria’s *R* identically benefits the prophage as well—no altruism is involved, merely an exact congruence of interests by both the viral prophage and its bacterial host. Thus, the virulence genes are of *indirect* value to the virus that carry them.

#### Gut Microbiomes—Habitat with High *R*

The guts of mammals (and presumably other groups too) are sites with concentrated nutrients and food for bacteria, likely increasing the bacterial *R* beyond that found in the environment while possibly leaving *b*_*o*_ relatively unaffected. As a result, a pattern emerges that parallels that of microbialization, namely, the bacteria of the gut are enriched with prophages, and moreover, those prophages produce virulence and pathogenicity. Specifically, in humans the gut bacteria, *Enterobacterales* and *Bacillales*, harbor significantly higher numbers of prophages and several members of Proteobacteria are the only ones enriched in prophages, compared with other taxonomic groups (López-Leal *et al*. 2022). Similar findings were previously reported for mice (Kim and Bae 2018). Moreover, López-Leal *et al*. 2022) also report that prophages are more abundant in bacteria associated with pathogenic phenotypes agreeing with previous reports, where clinical isolates of *Acinetobacter baumannii* and *Pseudomonas aeruginosa* showed a higher abundance of prophages than environmental isolates (Vallenet *et al*. 2008, Tsao *et al*. 2018). Thus, the *Acinetobacter* and *Pseudomonas* genera were enriched in prophages when these species displayed pathogenic phenotypes. Thus, the pattern of prophage enrichment and increased virulence in gut bacteria appears to share the same explanation as that for coral-reef microbialization because of the high *R* of bacteria in the gut environment.

#### Subsurface Ocean—Habitat with Low *R* and Even Lower *a*_*o*_ *b*_*o*_

Bacteria in the middle to deep ocean are enriched with lysogens in contrast to the predominantly lytic lifestyle of their photic-zone counterparts (Mizuno 2016, Luo *et al*. 2020, Yi *et al*. 2023 *cf*. Fig. 1e). Bacterial growth in the deep ocean is very slow with reported doubling times of 1.5 y (Gilgan *et al*. 2020). This pattern contrasts with the increased lysogeny in high-resource fast-growth situations such as eutrophic coral reefs and mammalian guts, as just discussed. Silveira *et al*. (2021) suggest that different mechanisms for coinfection account for these contrasting patterns. Here, an explanation based on the fitness switch, Eq. 17, must postulate that even though the bacterial *R* is lower in the deep ocean, the product, *a*_*o*_ *× b*_*o*_, is even lower so that condition B holds. This product may be lower because the micro turbulence at depths with limited wave action might lead to a very low *a*_*o*_.

#### Other Habitats With Increased Lysogeny

The prevalence of lysogeny increases with soil depth (Liang *et al*. 2020) and is high in the microbial mats of hot springs characterized by low mobility (Jarett *et al*. 2020).

Collectively, the habitat-lysogeny correlations constitute ecogeographic rules for the biogeography of virus that are logical counterparts of the many ecogeographic rules found in macro ecology. Classic macro ecogeographic rules include the pattern of lower clutch size on islands *vs* the mainland and many others (*cf*. Lomolino 2006, Gaston *et al*. 2008, Baiser *et al*. 2019).

### “Prophage Lock”

How long should prophages remain in bacterial genomes before switching back to produce lysis? The fitness switch, Eq. 17, suggests the prophage should remain within their hosts for a long time unless conditions substantially change.

Let the population growth for a bacterium infected with a prophage containing a virulence or other beneficial gene be *R*^*\**^, where *R*^*\**^ *> R*, the bacterial growth in the absence of the prophage. Condition B in Eq. 17 then has *R* replaced by *R*^*\**^. Because *R*^*\**^ *> R*, it is more difficult for the virus to switch back to the lytic condition, A, than if it were not carrying a beneficial gene. Thus, once a virus incorporates as a prophage containing a virulence or other gene benefiting the bacteria, it becomes locked into the lysogenic life cycle—it cannot revert back to the lytic life cycle without giving up the *R*^*\**^ it now enjoys. This situation is termed the “prophage lock.”

The possibility that prophages become locked into the lysogenic life cycle because they contain a gene that benefits their bacterial host may explain the longevity of prophages in host genomes through evolutionary time (Hedžet *et al*. 2022) as well as the domestication of prophages (Bobay *et al*. 2014). Indeed, spontaneous departure from the lysogenic state is rare, occurring at a frequency of less than once per million cell-generations (Aurell *et al*. 2002, Bæk *et al*. 2003).

Anthenelli *et al*. (2020) suggested that a virus incorporating into the bacterial host genome can be viewed as a “rachet” that they describe a process involving a reorganization of virus/bacteria community structure during microbialization. The prophage lock here is a wholly different idea that relies on the fitness advantage to a prophage of remaining in its host genome provided it carries a gene benefiting the host. Also, Sudhakari and Ramisetty (2023) have described the “grounding” of prophages in the host genome. They note that most putative prophages are relics of past phage/bacteria conflicts and consider them as “grounded” because they can no longer be excised from the bacterial genome. They interpret the grounding a result of phage/bacteria conflicts whereby the bacteria “win” by grounding the prophage in their genome to confer immunity to superinfection.

#### Virus as Vectors of Horizontal Gene Transfer

Virus can bring about horizontal gene transfer (HGT) (Canchaya *et al*. 2003, Gilbert and Cordaux 2017, Borodovich *et al*. 2022, Irwin *et al*. 2022). Prophage in one bacterial line can “pick up” a fragment of the host DNA that contains a gene, transition to the lytic phase, then infect other bacterial lines and ultimately transition back to a prophage in another bacterial line whereupon the foreign DNA is expressed there. The evolutionary question is why a virus should conveniently serve as a vehicle for transporting genes from one bacterial lineage to another.

The switch function, Eq. 17, suggests a scenario in which horizontal gene transfer makes evolutionary sense for the virus regardless of its effect on the bacteria. Suppose the bacterial environment is in continual fluctuation. If so, a high *R* supporting a lysogenic life cycle may fluctuate down to a low *R* favoring a lytic life cycle, whereupon the virus releases from the bacterial genome and lyses into the environment as virions. Next, these particles infect other bacteria. If *R* should fluctuate back up to a high value favoring a lysogenic life cycle, then the newly infecting virus incorporate into the genome of their new hosts and express their gene-containing fragment there.

This scenario derived from the model’s switch function predicts that horizontal gene transfer should proceed at a higher rate in fluctuating environments than in quiescent environments.

#### Three-Level (Tripartite) Holobionts

Holobionts consisting of three levels—virus, bacteria and eukaryotic host, often called “tripartite” holobionts, are receiving increased attention (Moran *et al*. 2005, Bordenstein *et al*. 2006, Kelly *et al*. 2014, Keen and Dantas 2018, Barr 2019, Belleghem *et al*. 2019, Bodner *et al*. 2021, Wahida *et al*. 2021, Zheng *et al*. 2023). A question raised by these contributions is whether a prophage gene expressed in a bacterium that itself resides in the microbiome of a eukaryotic host can improve the fitness of the eukaryotic host, producing a tripartite hologenome.

For viral-carried gene to affect the fitness of the host in a holobiont, a temperate virus must first be incorporated as a prophage into the bacterium—Eq. 17, Case B supplies the condition for this to occur. Next, the prophage-containing bacterium must be incorporated into the host’s microbiome. The process for entry into the host’s microbiome is host-orchestrated species sorting (HOSS, Roughgarden 2023). By this process, the bacteria must offer enough benefit to the host to avoid its exclusion by the host immune response. If a gene in the prophage of a bacterium in the host’s microbiome causes the bacterium to allocate sufficient resources to the host, then the host relaxes its immune screening of the bacterium and the prophage-containing bacterium becomes integrated in the host’s hologenome forming a three-level symbiosis. Thus, a gene in the prophage of a bacterium can theoretically contribute to holobiont fitness as a component of the hologenome provided the gene causes the bacterium it inhabits to contribute enough to the host fitness that it is allowed into the host’s microbiome—the result is a functionally integrated tripartite holobiont.

## Discussion

This paper introduces a simple model for the population dynamics of virulent phage and bacteria that features infection of bacteria by Poisson sampling of viral particles from the environment rather than mass-action collisions of virus with bacteria. Also, the new model is developed in discrete time with a time step equal to the viral latency period, rather than in continuous time and does not require assuming density-dependent bacterial population growth. The source of density dependence in the model lies in the kinetics of viral absorption onto the bacterial surface. The model of virulent-phage bacteria population dynamics was extended to temperate phage.

The condition for transition from a lytic to lysogenic phase was derived as a formula termed the “fitness switch”. The virus switches from lytic to lysogenic when its population grows faster with prophage than as virions produced by lysis of the infected cell. The switch function compares the product, base absorption probability *×* the per-virus burst size, with the bacterial population geometric growth factor, *R*.

The switch function is used to predict temperate viral-bacteria dynamics in several environmental scenarios: intervention that lowers probability of absorption, environmental deterioration that affects both virus and bacteria, the ecogeographic rule that environmental enrichment increases bacterial population growth (microbialization), fluctuating environmental conditions that promote horizontal gene transfer (HGT), and the integration of a prophage-containing bacteria population into the microbiome of a eukaryotic host that leads to a functionally integrated tripartite holobiont.

The key features of the model worth explicit experimental testing are the condition for increase when rare, Eq. 11, that is the criterion for where a lytic phage and bacteria can coexist and the extension of this condition as the fitness switch, Eq. 17, that is the basis for the model’s many qualitative predictions. The increase-when-rare condition would be tested with a lytic virus in an experimental setup wherein the virus *a*_*o*_ and *b*_*o*_ parameters were measured along with the bacterial *R*. If the *R* were experimentally raised, say by augmenting bacterial resources or changing environmental temperature, from a low value at which *a*_*o*_ *b*_*o*_ *> R* to a higher value at which *a*_*o*_ *b*_*o*_ *< R*, then the lytic virus should infect at low *R* and fail to infect at high *R*. The threshold *R* between success and failure to infect should occur at the experimental conditions where *R* satisfies *a*_*o*_ *b*_*o*_ = *R*.

The fitness switch condition would be similarly tested with a temperate virus. If the *R* were experimentally raised from a low value then the prediction is that the virus would integrate until experimental conditions pass the threshold at which *a*_*o*_ *b*_*o*_ = *R*. That is, initially when *a*_*o*_ *b*_*o*_ *> R*, virus added to a bacteria population should infect and then lyse whereas when *a*_*o*_ *b*_*o*_ *< R* the introduced virus should infect and then integrate. The integration is presumably measured by noting that the concentration of viral integrase enzymes suddenly rises from a low level to a high level as the threshold is passed. Reversing this sequence should induce the prophage to initiate lysis. Undoubtedly experimentalists can suggest better setups than the author.

Another feature needing explicit test is the model’s assumption of density-dependent absorption. Studies of viral absorption kinetics like those cited earlier in the paper should be conducted in the future at a full range of multiplicities of infection (MOI).

The model in this paper falls into the category of dynamical models with implicit resources as contrasted with models providing explicit resources as reviewed, for example, in Weitz (2015, pp. 58–74). One circumstance in which an explicit-resource model might be preferred is where resource-limited bacterial growth might account for a decline in the physiological state within the bacteria as a carbon source becomes depleted—this in turn might affect viral growth within the bacterial hosts (Golec *et al*. 2014, Attrill *et al*. 2023).

The model can be extended to account for additional phenomena. Notable among these is the evolution of phage resistance by the bacteria. The rise of antibiotic drug resistance in bacteria is fueling interest in phage therapy as a possible way to kill these pathogenic bacteria. But the prospects for using phages as biological control are tempered by the ability of the bacteria to evolve resistance to infection by phages (Azam and Tanji 2019, Monteiro *et al*. 2019, Torres-Barceló *et al*. 2022). A future paper building on this paper’s model will explore theoretically the coevolution of phage with bacteria and specifically, the evolution of phage resistance.

This paper’s model is relevant to hypotheses in the virus ecology literature about how viruses influence microbial community structure. Thingstad (Thingstad and Lignell 1997, Thingstad 2000, Winter *et al*. 2020) advanced a hypothesis known as “Kill the Winner” (KtW). The idea is that a bacteria population that happens to be abundant is, as a result of its abundance, subject to more infection from lytic viruses than less abundant bacteria. Thingstad refers to this effect as “negative frequency selection” and explicitly cites the Lotka-Volterra predator-prey equations as the starting point for a KtW model. That is, when the prey abundance increases, the predator abundance increases soon thereafter, chopping down a temporary excess of prey. When extended to multiple species of bacteria, the hypothesis is that whenever one bacterial strain increases, perhaps because of being a better competitor, its success is countered by increased viral infection, thereby preventing any one bacterial strain from dominating. This process is hypothesized to maintain microbial diversity. Lytic virus are thus envisioned as keystone predators.

Lytic viruses that control an algal bloom in the way envisioned under KtW were documented by Bratbak *et al*. (1993) who report that blooms of a marine coccolithophorid off the coast of Norway were in some cases succeeded by an increase in the abundance of viruses. They further report that viral lysis could account for 25 to 100% of the net mortality of the coccolithophorids and that by the time the blooms declined the concentration of free algal viruses was seen to have increased. However, Evans and Brussaard (2012) report that in the Southern Ocean both viral lysis and microzooplankton grazing together control cyanobacterial populations, whereas viral control of eukaryotic algae was comparatively minor. And finally, Needham *et al*. (2013) report that off the coast of California near Santa Catalina Island common bacterial and viral taxa were consistently dominant and that “relatively few displayed dramatic increases/decreases or ‘boom/bust’ patterns that might be expected from dynamic predator-prey interactions.” Thus, empirical support for the picture envisioned in the KtW hypothesis is mixed.

The model in this paper is not consistent with a KtW picture based on the Lotka-Volterra predator-prey model because in this paper’s model, a lytic virus population comes to equilibrium with its bacterial host. In this paper’s model, the virus and bacteria do not oscillate as they do in a Lotka-Volterra-like model such as Campbell (1961). Hence, in this paper’s model the virus population does not chase abundant bacteria chopping down their abundance and killing the winner. Instead, in this paper’s model, the lytic virus coexists with the winner even though this may result in the exclusion of the losers and an accompanying loss of microbial diversity.

Furthermore, according to the KtW picture, a high abundance of winning bacterial strains should soon be followed by a high abundance of virus that are in the process of killing the winners, resulting in a high virus to microbe ratio (VMR) in the environment. However, Forest Rohwer and colleagues (Knowles *et al*. 2016) demonstrated across multiple ecosystems and laboratory situations that virus are relatively less abundant in the environment when occurring with high bacterial densities. Specifically, microbialization on coral reefs along with their elevated bacterial abundances is associated with low VMR rather than high VMR as expected by the KtW hypothesis.

As a result, Rohwer and colleagues (Knowles *et al*. 2016, Silveira and Rohwer 2016) proposed an alternative hypothesis they termed “Piggyback the Winner” (PtW). By this hypothesis, virus become lysogenic in conditions of high bacterial abundance resulting in few virus particles being found outside the bacterial cells because they are residing inside the bacteria as prophages, leading to a low VMR.

On its face, the fitness switch in this paper supports the PtW hypothesis because, as mentioned earlier, the higher *R* in the nutrient-rich eutrophic conditions that cause microbialization leads to satisfying Condition B in Eq. 17, indicating a switch from lysis to lysogeny.

In contrast to the fitness-switch theory espoused in this paper, Silveira and Rohwer (2016) propose a hypothesis for the increase in lysogeny during microbialization based on the observation that mucosal surfaces in microbialized environments facilitate viral colonization of bacteria (Barr *et al*. 2015). However, this increases the absorption parameter, *a*_*o*_ in Eq. 17, favoring the lytic phase according to Condition A rather than the lysogenic phase according to Condition B. So, according to the fitness switch model, an increased rate of phage-bacteria encounters in mucus cannot explain the increase in lysogeny during microbialization because the argument goes the wrong way.

Alternatively, Luque and Silveira (2020) also note that the high viral densities in microbialized situations leads to increased coinfections which is known to promote lysogeny, as discussed earlier. This might contribute to favoring lysogeny. However, increasing DOM increases the bacterial *R* and can flip the fitness switch from lytic to lysogeny by itself. Therefore, this paper is consistent with the PtW hypothesis if understood as follows: the reason for PtW is the high *R* associated with eutrophication, although a role for increased coinfection in eutrophic conditions cannot be ruled out.

KtW and PtW need not necessarily be viewed as competing hypotheses. Indeed, Chen *et al*. (2021) argue that KtW and PtW “are not mutually exclusive and need to be combined.” Alternatively, one might simply stipulate that KtW pertains only to lytic viruses whereas PtW pertains only to temperate viruses. With this stipulation, whether KtW is correct for lytic virus can be examined on its own merits regardless of whether PtW is also correct for temperate virus, and *vice versa*.

## Acknowledgments

The author thanks Forest Rohwer, Toni Luque, Sergio Cobo-Lopez, James Nulton and Anca Segall of the SDSU BioMath Group, Seth Bordenstein, Sarah Bordenstein, Andrew Read, Ottar Bjørnstad and members of the Penn State Microbiome Center, Jaime G. Lopez of the Stanford Medical School and Richard White III of UNC Charlotte for discussion about this paper. The author also thanks two anonymous reviewers for helpful comments. This article is Contribution #2 from a project, The Theory of Holobiont Evolution, funded by the Gordon and Betty Moore Foundation through Grant GBMF10000 to the University of Hawaii.

## Supplementary Material

### Terminology

A virus particle that infects a bacterium is a “bacteriophage” (eats bacteria) or just “phage”, for short. A complete virus particle is a “virion”—it has a protein coat that is the “capsid” enclosing its nucleic acid genome.

A “virulent phage” enters a bacterial cell, commandeers its replication machinery and after a “latency period”, releases copies of itself into the environment, killing the bacterium. “Lysis”, from the Latin word for “a loosening”, is the process by which the copies are released. The set of released copies is the “burst”, the number of copies within the burst is the “burst size”.

A “temperate phage” may either reproduce new virus particles for release into the environment while killing the bacterium *or* may integrate into the bacterium’s genome and reproduce jointly with it. In the former case, the virus, like a virulent virus, commandeers the bacterium’s replication machinery to produce copies of itself that after a latency period lyse from the bacterium killing it. This sequence is the “lytic pathway.” In the later case, the virus integrates into the bacterial genome becoming a “prophage” that replicates along with the bacterial genome as the bacterium reproduces. The bacterium is not killed. A bacterium containing a prophage within it is a “lysogen”. This sequence is the “lysogenic” pathway.

Following incorporation into a bacterium’s genome, a temperate phage may either remain incorporated as a prophage during subsequent bacterial cell divisions. *Or*, it may become virulent by de-incorporating from the bacterium’s genome and proceed to produce new virus particles by lysis into the environment while killing the bacterium it formerly resided in.

A prophage may augment a bacterium whose genome it resides in by providing the bacterium with a novel gene expressed through a process of “lysogen conversion” *i.e*., the bacterium is converted from lacking a function into expressing a function. Prophages often encode genes called “morons” that are not directly involved in viral replication and can confer a benefit to their bacterial host. Such genes are independent transcriptional units of DNA that are expressed while the phage is in the prophage state. Morons can enhance the virulence of bacteria to their own hosts by supplying phage-encoded toxins.

Terms in virology are defined relative to experimental protocols that have no ready counterpart in ecology, hindering the translation of terms between these disciplines. Virologists define the “multiplicity of infection” or MOI with a protocol whereby phage are added to bacteria in a medium. The ratio of the number of phage to the number of bacteria initially added to the medium is the MOI. In this paper, *v*(*t*) = *V*(*t*)/*N*(*t*), is the ratio of virus particles, *V*(*t*), to bacteria, *N*(*t*), in their source pools at time, *t*. Hence, the MOI roughly corresponds to *v*(0) in this paper.

A “one-step growth curve” is a curve of viral abundance in continuous time from the time of absorption until the time of lysis. The one-step curve may include information about the abundance of virus both within the bacteria as well as in the media. A “multi-step growth curve” is an experimental iteration of the one-step curve obtained by allowing the experiment to run beyond the first step. In this paper, viral abundance, *V*(*t*), and bacterial abundance, *N*(*t*), are curves in discrete time and are recorded solely at the end of each latency period, not within the latency period. They refer to abundance in the environmental source pools, not to virus within bacteria during a time step. Furthermore, Gadagkar and Gopinathan (1980) define an “effective multiplicity of infection” as the “mean number of phages adsorbed per infected bacterium”. They also define the “effective burst size” as the “mean number of phages liberated per phage adsorbed.” In this paper, at each time step, *m* indexes the number of virus in a bacterium where *b*(*m*) is the burst size from a bacterium infected with *m* virus, and although not used here, *b*(*m*)/*m*, would represent the effective burst size.

### Campbell Model of Virulent Phage/Bacteria Population Dynamics

In 1961, Allan Campbell proposed a mathematical model for the population dynamics of virulent phage and bacteria. The Campbell model (Campbell 1961) introduced features that have persisted in phage/bacteria models until the present day. The Campbell model is:

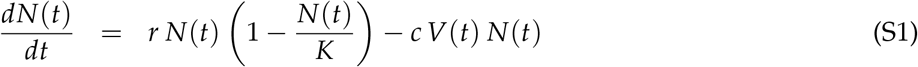

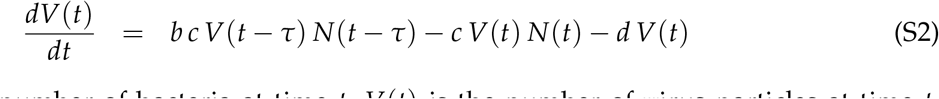

where *N*(*t*) is the number of bacteria at time *t, V*(*t*) is the number of virus particles at time *t, N*(*t − τ*) is the number of bacteria at time *t − τ, V*(*t − τ*) is the number of virus particles at time *t − τ, τ* is a time lag, *r* is the bacterial intrinsic rate of increase, *K* is the bacterial carrying capacity within its environment, *c* is the rate of conversion of mass-action collisions between virus particles and bacteria into infections, *b* is the burst size, and *d* is the per capita disintegration rate of virus particles. (The Campbell model was originally written for a phage/bacteria system in a chemostat. Here the chemostat nutrient flow terms are omitted.)

The Campbell model is a pair of nonlinear delay-differential equations. The first term of Eq. S1 assumes the bacteria grow logistically in the absence of virus. The second term is the loss of bacteria from infection brought about by mass-action random collision of bacteria with virus particles. The first term in Eq. S2 is the production of new virus particles that depend on the collisions that took place *τ* units of time previously. The time lag, *τ*, called the latency, is the time needed for the infecting virus particles to manufacture and release new virus particles using the bacteria’s nucleotide replication machinery. The next term indicates the loss of free virus particles from the infections taking place at the present time, *t*. Thus, the virus particles disappear for the duration of the latency period. After infecting bacteria at time *t − τ* they reappear at time *t* as newly minted virus particles.

The Campbell model treats the bacteria and virus as *two* separate populations. This type of model is distinguished from those used in epidemiology that pertain to a *single* population containing multiple classes (Kermack and McKendrick 1927, Anderson and May 1979, Keeling and Rohani 2008, Diekmann *et al*. 2013, Bjørnstad 2018.) The time units in single-population models pertain to the time course of the disease in the host, say days to months, whereas the time units in the Campbell model relate to the kinetics of bacterial and viral replication, say minutes. Some models have been proposed for marine virus and also for bacteria in chemostats that are hybrids between the Campbell-type population model and an epidemiology type model (Beretta and Kuang 1998, 2001, Bull *et al*. 2006).

#### Equilibrium Coexistence of Virus and Bacteria

The equilibrium point of virus-bacteria coexistence in the Campbell model, 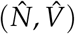, is found by setting 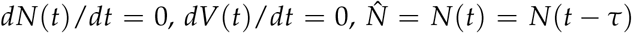, and 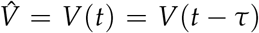 in Eqs. 1 and 2 from the main text yielding

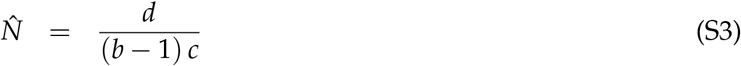

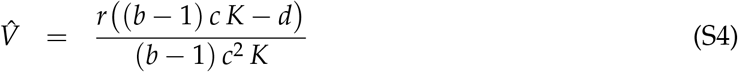

The equilibrium point is independent of the latency time, *τ*, although *τ* does influence the stability of the equilibrium. The condition for a feasible equilibrium point of coexistence between virus and microbe is read off from the numerator of Eq. S2 as

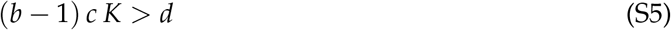

This condition means the birth of virus has to exceed the death of virus. This condition can always be satisfied if *K* is large enough. Specifically, the bacterial carrying capacity, *K*, must exceed

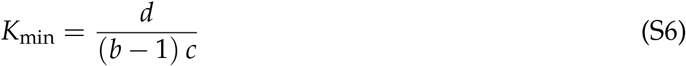

for the virus and bacterial populations to have positive equilibrium values. In fact, *K*_min_ works out to be the same as 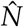. Therefore, *K* must merely be greater than the bacterial abundance expected in the presence of the virus for the equilibrium in Eqs. S1 and S2 to be positive. The 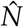 is independent of *r* and *K*, indicating that its abundance is controlled by the virus’ parameters. The 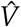 does depend on *r* and *K*, and increases with the bacteria’s *r* and decreases with the bacteria’s *K*. This dependence of the bacterial abundance on the properties of the virus and of virus abundance on properties of the bacteria is a well known property of Volterra-style predatorprey models in ecology (*c.f*. Roughgarden, 1998, p. 266 *ff*).

**Table S1:** Example Parameters in Campbell Model

|  |  |
| --- | --- |
| $r$ | 0.0347 |
| $K$ | $10^8$ |
| $\tau$ | 15 |
| $b$ | 40 |
| $c$ | $10^{-4}$ |
| $d$ | 0.693 |

#### Parameters for Campbell Model

Order-of-magnitude guesses at parameter values for the model may be constructed from the literature. Consider the parameters for bacterial logistic growth, *r* and *K*. In the laboratory, the doubling time of *E. coli* is 20 minutes although it can be much slower in the wild (Gibson *et. al*. 2018). Assuming time units of minutes, this laboratory doubling time corresponds to *r* = 0.0347 (*i.e*. 0.0347^20^ = 2.00). Next, in the human gut, which is the habitat of *E. coli*, this bacterium is typically present in 10^6^ colony forming units per gram of stool (Foster-Nyarko and Pallen 2022). A typical value for the amount of faeces in a human is one lb or 450 grams (Alexa Answers, Amazon, retrieved July 21, 2022). So, a possible value for *K* is 10^8^.

The time between the viral adsorption and first appearance of viral progeny is 10–15 minutes (Endy *et al*. 1997, Purohit *et al*. 2005). So, a possible value for the latency, *τ*, is 15 minutes.

The burst size from microbes infected with about two virus is about 80 virus particles (Gadagkar and Gopinathan 1980, Figure 4), so *b* will be taken as 40.

About one in 10^3^–10^4^ collisions between a virus and bacterium lead to a specific binding between the cellular receptor and the virus attachment protein on the surface of the virus (Philipson 1983). So in the model *c* is taken as 10^*−*4^.

The half-life of free infectious influenza A virus is *≈* 3 hours (Baccam et al 2006). If the half life is taken on the order of 1 hour, the disintegration rate of free virus, *d*, works out to be about 0.693 (*i.e. d* = *−* ln(1/2)).

**Figure S1:**
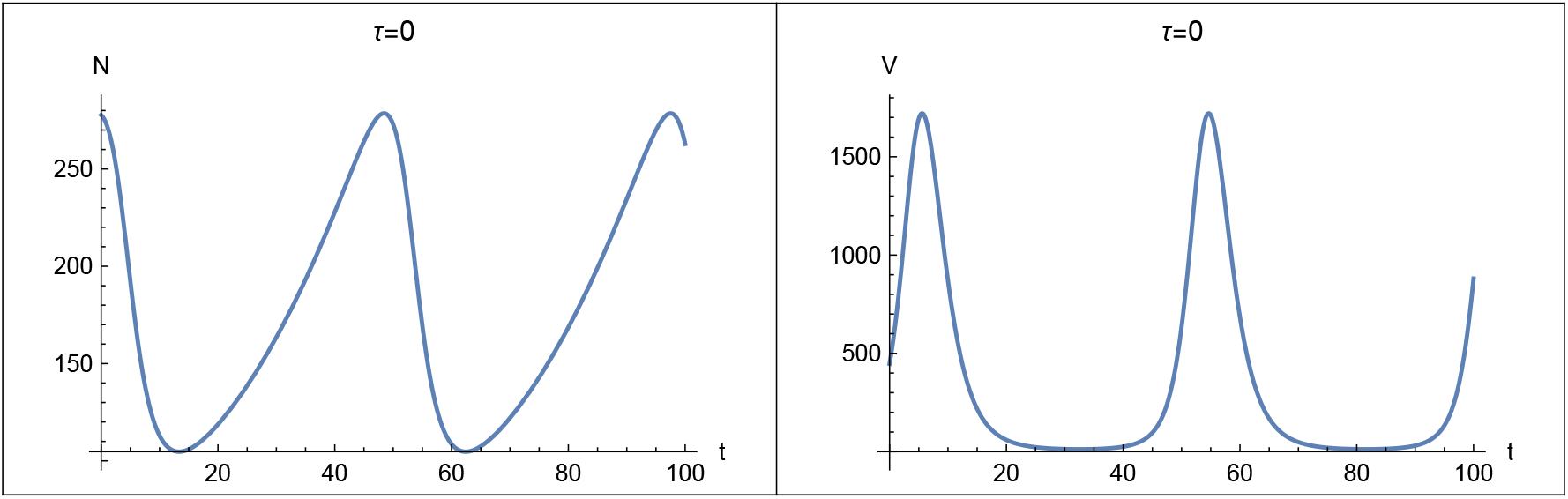
Virus-Bacteria oscillation from the Campbell model with parameters from Table S1 excepting τ = 0. Left is number of bacteria vs. time and right is number of virus vs. time. Initial conditions are 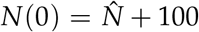 and 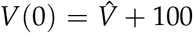. The period of the oscillation is about 55 minutes. Integration is from t = 0 to 100 minutes.

These guesses at plausible parameter values are summarized in Table S1.

Guesses at plausible parameter values are summarized in Table S1. Using these parameters, the equilibrium from Eqs. S1 and S2 at which bacteria and virulent virus coexist out to be

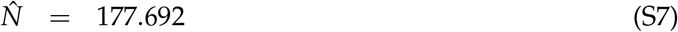

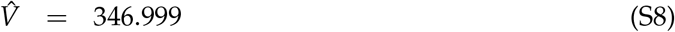

#### Stability of Equilibrium with No Latency, *τ* = 0

The coexistence of bacteria and a virulent virus require both that the equilibrium solutions for 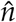 and 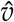 from Eqs. S1 and S2 be positive *and* that the equilibrium point be stable to perturbations.

To begin, assume that the latency, *τ*, is 0, indicating an instantaneous conversion of virus infections into newly released virus particles. This assumption converts the Campbell model from a pair of delay-differential equations into a pair of Volterra-style ordinary differential equations that can be analyzed with conventional methods. The details of the analysis are recorded in the Supplementary Material.

With *τ* = 0, the equilibrium is always stable. The trajectories reveal a virus-bacteria oscillation through time that winds in to the equilibrium point. The period of the oscillation is determined by *d* and *r*, independent of *b, c* and *K*. The oscillation speeds up as *d* and/or *r* are increased. A high *K* diminishes the stability of the virus-bacterial equilibrium point.

The Campbell model, Eqs. 1 and 2, with *τ* = 0 can be integrated with the NDSolve[] function in *Mathematica*. The solutions appear in the time domain as a nearly endless virus-bacteria oscillation as illustrated in Figure S1 and in a parametric plot as a nested set of loops, each representing a different initial condition, as illustrated in Figure S2. If the bacterial carrying capacity within the host is arbitrary set lower than 10^8^, say approaching *K*_min_ from Eq. S4, then the equilibrium becomes increasingly stable and the trajectories more rapidly spiral into the equilibrium point (not illustrated).

**Figure S2:**
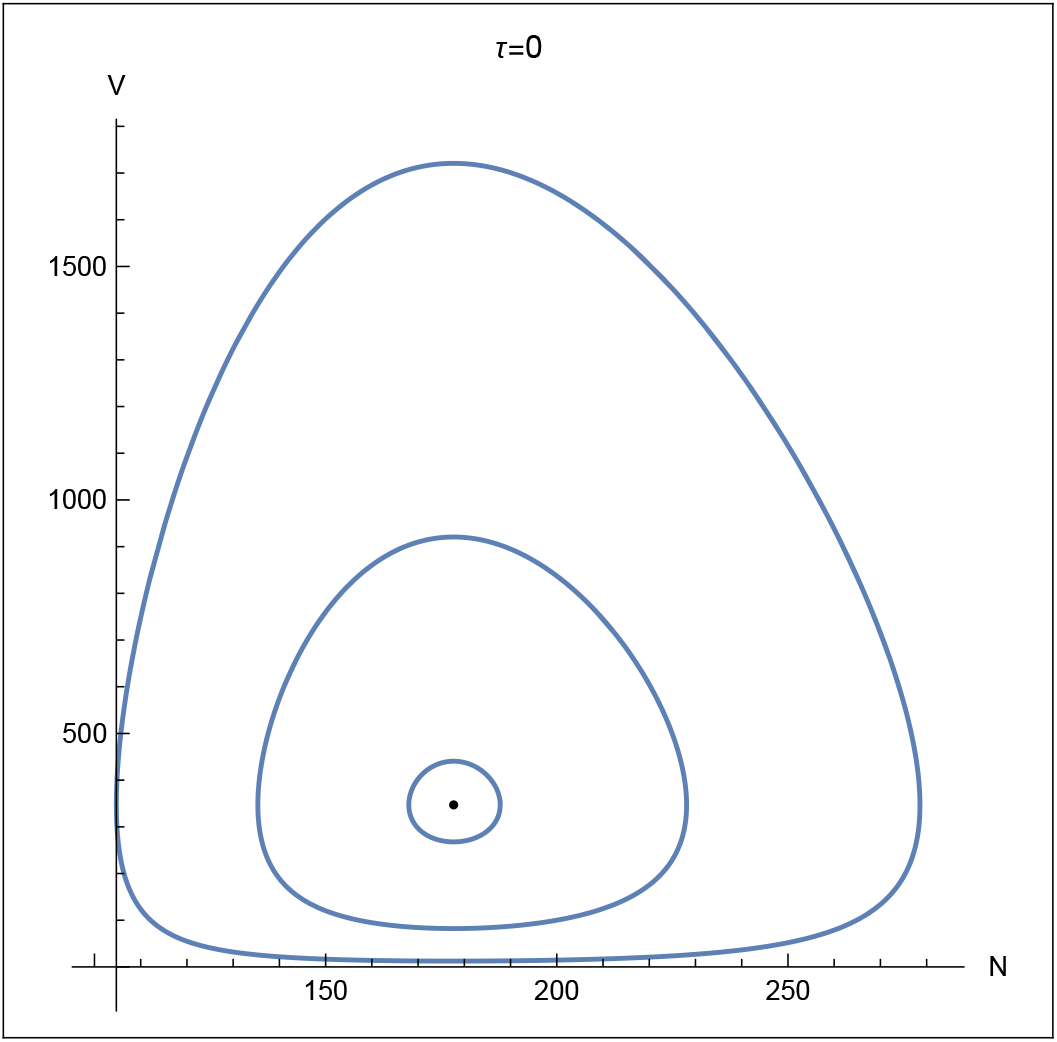
Parametric plot of the Campbell model with parameters from Table S1 excepting τ = 0. Horizontal axis is number of bacteria, vertical axis is number of virus, dot in center is the equilibrium point, 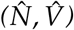. Outer loop was started at 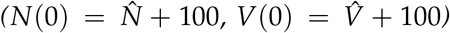, middle loop started at 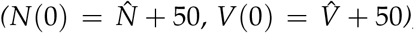, and inner loop started at 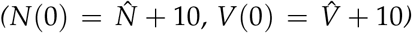. Integration is from t = 0 to 100 minutes.

#### Stability Analysis of Campbell Model with *τ* = 0

The Jacobian matrix, *J*, with *τ* = 0 evaluated at the equilibrium yields

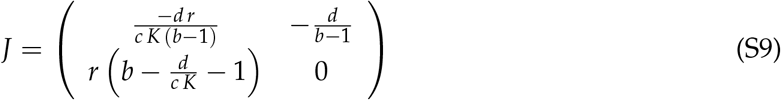

The eigenvalues of *J, λ*_1_ and *λ*_2_ are a pair of conjugate complex numbers the real part of which is

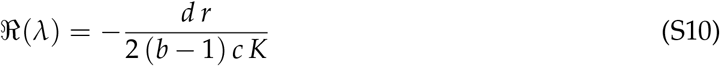

Because ℛ(*λ*) is always negative, the equilibrium is always stable. And because the eigenvalues are complex, the trajectories reveal a virus-bacteria oscillation through time that winds in to the equilibrium point in a flow pattern representing a “stable focus”. The period of the oscillation is approximately

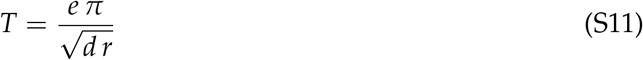

The period is determined by *d* and *r*, independent of *b, c* and *K*. The oscillation speeds up as *d* and/or *r* are increased.

**Figure S3:**
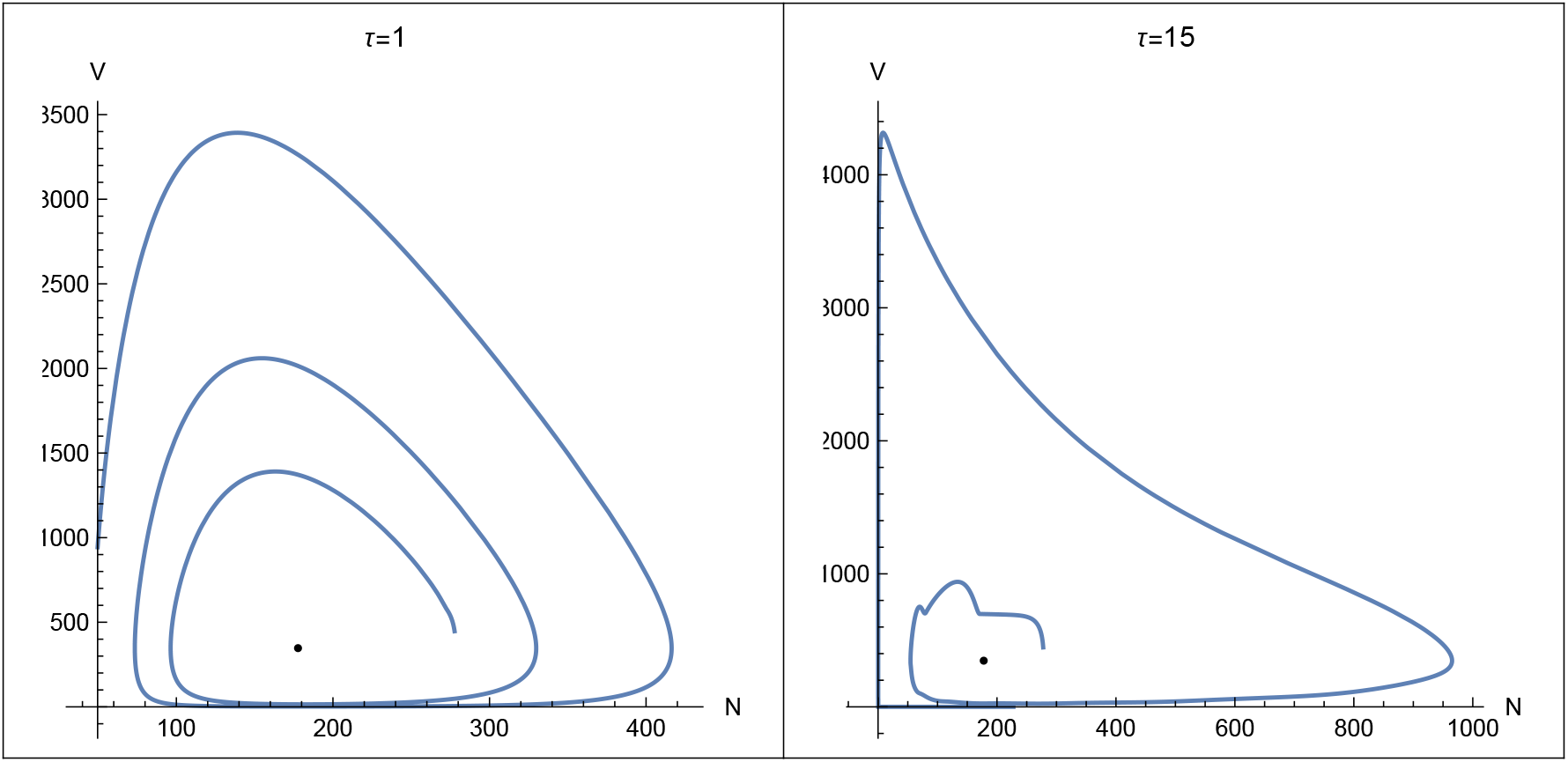
Left: Parametric plot of the Campbell model with parameters from Table S1 excepting latency, τ = 1 minute. Right: Parametric plot of the Campbell model with parameters from Table S1. Horizontal axis is number of bacteria, vertical axis is number of virus, dot in center is the equilibrium point, 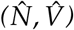. Initial condition is 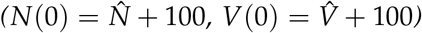. Integration is from t = 0 to 150 minutes on the left and to 400 minutes on the right.

Even though ℛ(*λ*) is always negative, its magnitude varies inversely with the bacterial carrying capacity, *K*. Hence, a high *K* diminishes the stability of the virus-bacterial equilibrium point. Indeed, if *K* is infinite, the equilibrium point is neutrally stable and the flow of trajectories represents a “center” as occurs in the original Volterra predator-prey model (Volterra 1926; for a summary *cf*. https://en.wikipedia.org/wiki/Lotka-Volterra equations, retrieved July 28, 2022).

With the parameters of Table S1,

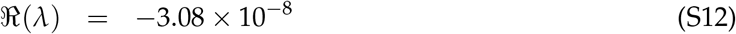

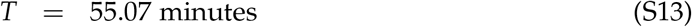

The ℛ(*λ*) for these eigenvalues is nearly zero because *k* is 10^8^ in Table S1 and thus the equilibrium is, practically speaking, neutrally stable.

#### Stability of Equilibrium with Latency, *τ >* 0

The introduction of the latency time lag, *τ*, destabilizes the equilibrium at which virus and bacteria coexist. Although stability conditions for a system of delay-differential equations cannot generally be analyzed mathematically, the NDSolve[] function in *Mathematica* can successfully numerically integrate the Campbell model with a fixed time lag, *τ*. The left side of Figure S3 shows the parametric plot of virus and bacteria through 150 minutes with a small latency, *τ* = 1 minute. The virus and bacteria populations were started toward the right of the equilibrium point, at 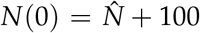 and 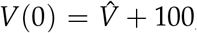, respectively. The trajectories wind out from the initial condition, spiraling ever farther from the equilibrium point.

The dynamic instability caused by the latency period becomes even worse with the realistic latency period of *τ* = 15 minutes, as was recorded in Table S1. The right side of Figure S3 shows the parametric plot of virus and bacteria through 400 minutes for a 15 minute latency. The virus and bacteria populations were again started toward the right of the equilibrium point, at 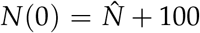 and 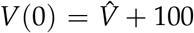, respectively. The trajectories show a jagged start because the initial condition for a delay-differential equation should specify the value of the variables at time *t* = 0 *and* at all times in the past until *t* = *−τ*. Because the value of the variable is unknown before *t* = 0, the value at *t* = 0 is attributed to the earlier times. After a while, the trajectory catches up with its history and the trajectory smooths out. Thereafter the trajectories wind out and quickly hug the axes.

Campbell in 1961 may not have been aware of the destabilizing role of the viral latency period. At that time he wrote “If a virulent phage and a susceptible bacterium are mixed in an open growth system, such as a chemostat, one expects that, in general, the concentration of the two will approach some stable steady state value.” (Campbell 1961, p157–8). But twenty years later, in 1981, he wrote that “Stable coexistence of phage and bacteria in chemostats has been verified experimentally. Comparison of theory with experiment revealed one discrepancy: Although the concentrations of bacteria and phage stabilized at approximately the predicted values, theoretical analysis and computer simulations predicted unstable oscillation rather than a stable steady state.” (Campbell 1981, p. 61). Indeed, researchers testing a version of the Campbell model with chemostats in 1977 wrote, “the time lag due to the latent period of the phage has a destabilizing effect which may be stronger than the opposite, stabilizing, effect of [bacterial population] regulation (Levin *et. al*. 1977, p. 11).

#### Critique of Campbell Model

Fifty-six years after the ground-breaking Campbell (1961) model appeared, Krysiak-Baltyn *et al*. (2017) published an extensive review of models for virulent phage and bacteria. They observed that “The basic model initially suggested by Campbell, and later refined by Levin and colleagues [Levin *et al*. 1977], has remained in use since first publication. Later studies have mostly made minor adjustments to the existing model to study dynamic behavior.” (p. 964) They further observed that “Earlier studies on simple chemostat models indicated that the dynamics of the model are not in good quantitative agreement with experimental data. In particular, the stability of the simulated models is lower than expected, with many exhibiting heavy oscillations and extinction events where stable steady-states exist experimentally.” (p. 964) Krysiak-Baltyn *et al*. (2017) conclude that “On a practical level, the usefulness of computational models of bacteria and phages has not yet been established. The simpler chemostat models may first need to become more accurate” (p. 965).

Thus, the Campbell (1961) model suffers quantitative limitations. First, it predicts instabilities to virulent virus/bacteria population dynamics that are contradicted experimentally. Second, the Campbell (1961) is so mathematically challenging that its use is impractical in biology. How to mathematically analyze a system of nonlinear delay-differential equations is an ongoing subject of research in applied mathematics (Tang and Zou 2008, Huang et. al. 2010; indeed many investigators simply put the time lag, *τ*, equal to zero to evade the problem, converting the delay differential equations into ordinary differential equations). And as discussed in the text of the main document, the qualitative picture provided by the Campbell model is problematic. The Campbell model envisions that virus/bacteria population dynamics involves two counteracting forces—the stabilizing force of density-dependent bacterial population growth and the destabilizing force of time lags arising from the virus latency period. The outcome is a balance of these forces. This qualitative picture might not be accurate.

### Density-Dependent Burst Size

Several choices are available for the burst size from a bacterium as a function of the number of virus in it, *b*(*m*). The burst size might increase linearly with *m* so that *b*(*m*) = *b*_*o*_ *m* where *b*_*o*_ is the contribution to the overall burst made by a single viral particle (Patel and Rao 1984, Figures 1–2). This case is for no density dependence. Alternatively, *b*(*m*) might be a constant, *b*(*m*) = *b*_*o*_, indicating that the burst size does not increase for *m* beyond the initial infection. A constant *b*(*m*) indicates strong density dependence and is consistent with surveys reporting that burst sizes are characteristic of various viral taxa (Parada *et al*. 2006, Ranasinghe 2019). Another possibility is to assume *b*(*m*) is a convex increasing function of *m* (Gadagkar and Gopinathan 1980, Figures 4–5). In this case, the burst-size function might be taken as *b*(*m*) = *b*_*o*_ *m*^*α*^ where *α* is an exponent between 0 and 1, say 1/2. A final possibility is to allow *α* to be negative, say *α* = *−*1, to indicate that burst size varies inversely with the multiplicity of infection, indicating extremely strong density dependence as a response to “superinfection” (Brown and Bidle 2014). All these possible relations between burst size and *m* may be subsumed by taking *b*(*m*) = *b*_0_ *m*^*α*^ with *α* as 1, 1/2, 0, or -1. Here, the cases of *α* = 1 and *α* = 0 will be investigated analytically. Results (not shown) for other *α* are similar.

**Figure S4:**
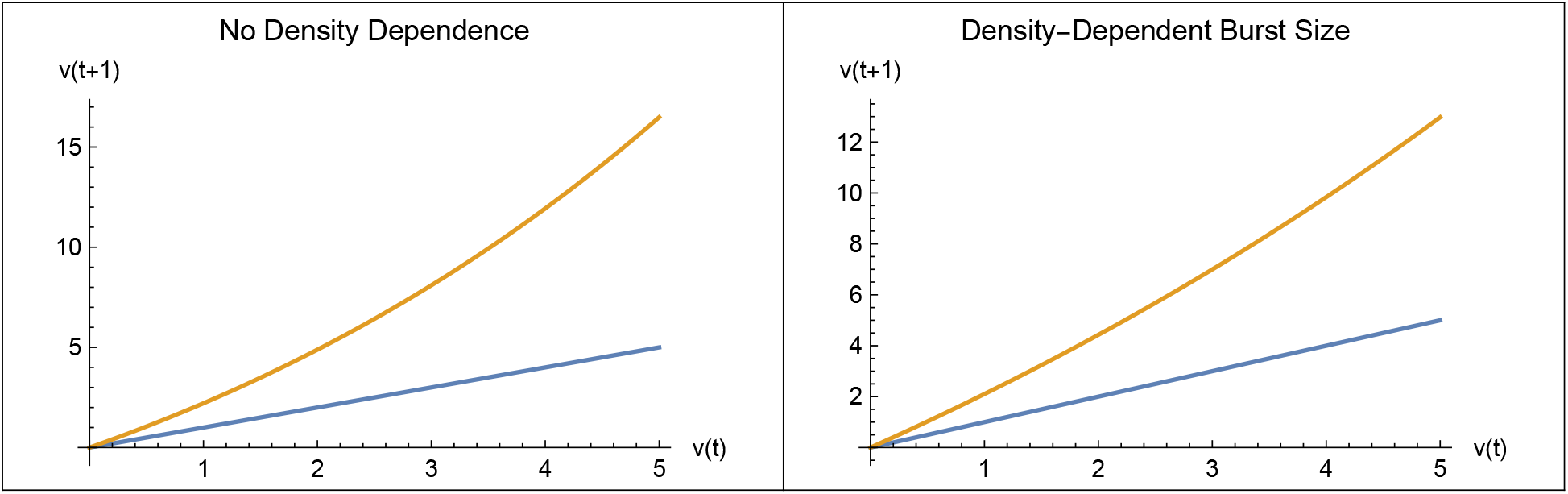
v(t + 1) vs. v(t), in brown, for no density dependence (left) and for a density-dependent burst size (right) from Eqs. 10 and 13, respectively. The diagonal line, in blue, indicates v(t + 1) = v(t). An intersection of the brown curve with the blue line would indicate an equilibrium point where virus and bacteria coexist, provided the equilibrium is stable. Because the curve does not intersect the line in either plot, no coexistence occurs in either case. Here, a_o_ = 0.1, b_o_ = 40, and R = 2

**No Density Dependence:** *a*(*v*(*t*)) = *a*_*o*_, *b*(*m*) = *b*_*o*_ *m*

Here,

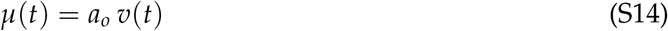

and, upon doing the summation in Eq. 4,

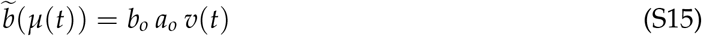

**Figure S5:**
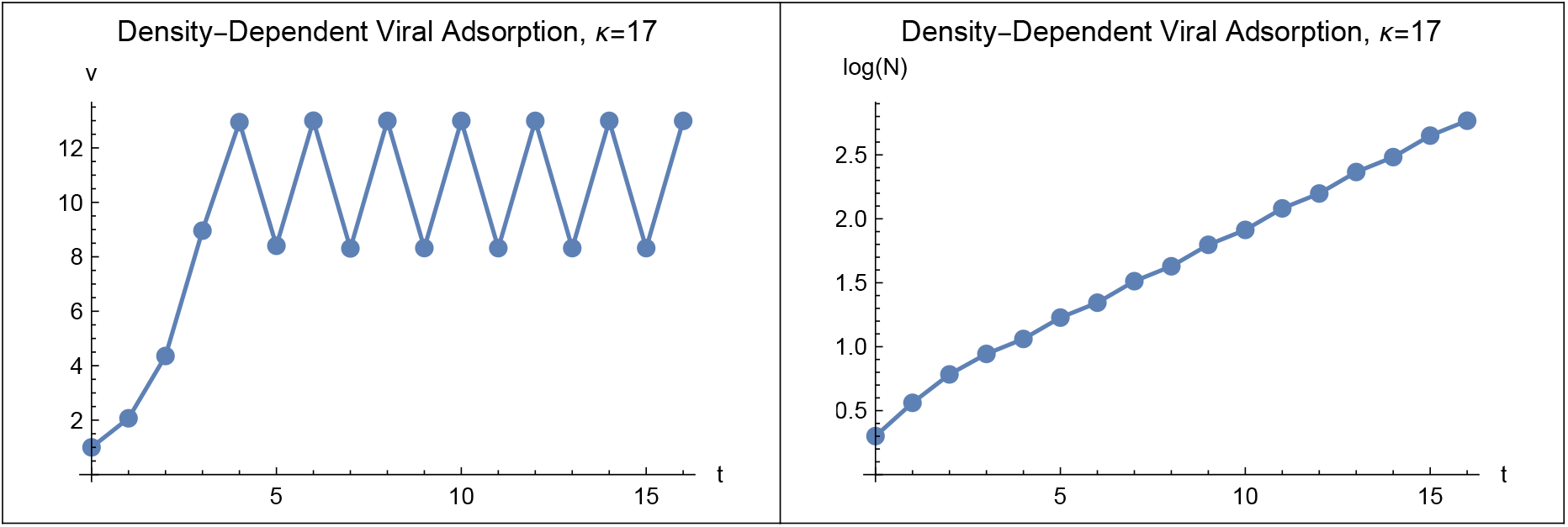
Left: Trajectory of v(t) showing a two-point limit cycle around the unstable equilibrium at 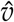. Right: Trajectory of log_10_(N(t)) showing a two-point oscillation in the exponential bacterial population growth. Here, κ = 12, v(0) = 1, N(0) = 2, t_max_ = 16, κ = 12, a_o_ = 0.1, b_o_ = 40 R = 2 and t_max_ = 16.

so Eq. 5 becomes

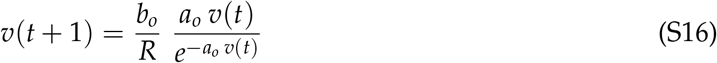

Analysis of this case appears in Figure S4 (Left).

**Density-Dependent Burst Size:** *a*(*v*(*t*)) = *a*_*o*_, *b*(*m*) = *b*_*o*_

Here,

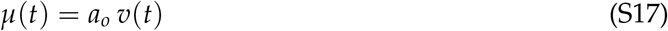

and, upon doing the summation in Eq. 4,

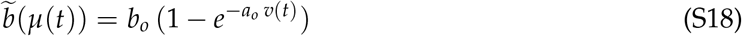

so Eq. 5 becomes

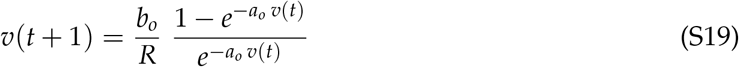

Analysis of this case appears in Figure S4 (Right).

A point of possible coexistence occurs at the value of *v* for which *v*(*t* + 1) equals *v*(*t*). This condition is shown graphically in Figure S4. An intersection of the curve for *v*(*t* + 1) with the line for *v*(*t* + 1) = *v*(*t*) would indicate an equilibrium point for possible virus/bacteria coexistence. The curve and line do not intersect in either case. Therefore coexistence, not surprisingly, is impossible without any density dependence, but surprisingly, is also impossible if the density dependence is solely in the burst size assuming the function relating burst size to density is *b*(*m*) = *b*_0_ *m*^*α*^. (Illustrated only for *α* = 0, 1.)

### Trajectories with *κ >* 14

Figure S5 illustrates trajectories of both virus and bacteria for *κ >* 14. Trajectories illustrate a two-point limit cycle of *v*(*t*) around the unstable 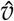. The trajectory of *N*(*t*) also shows a two-point oscillation around the asymptotic growth factor although again, the scale on the graph makes the oscillations difficult to see.

The existence of a two-point limit cycle around the unstable equilibrium in Figure S5 is consistent with known dynamics for the discrete-time logistic equation (May 1976.) The possibility of more complex oscillations and chaotic trajectories with *λ*_+_ *< −*1 remains to be explored. For the purposes of this paper, a comparison of the oscillations for this model in Figure S5 with those in Figures S1–S3 shows that the oscillations in the present model are bounded from above and below whereas those in the Campbell (1961) model approach arbitrarily close to the axes, implying population extinction.

### Poisson Summation Identities

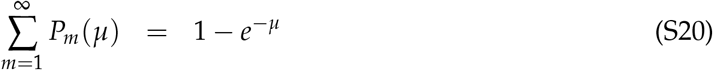

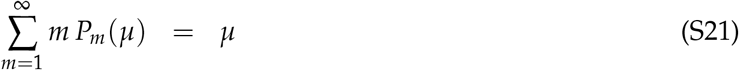

## Footnotes

3 *cf*. Poisson summation identities in Supplementary Material

